# Modeling host-microbe interactions in immunocompetent engineered human gut tissues

**DOI:** 10.1101/2025.05.02.651468

**Authors:** Rubén López-Sandoval, Marius F. Harter, Qianhui Yu, Laura Gaspa-Toneu, Irineja Cubela, Kristina Kromer, Julien Aubert, Adrian M. Filip, Lukas Kaltenbach, Marina Almato-Bellavista, Ninouk Akkerman, Yannik Bollen, Salem Munteanu, Justine Fidelin, Angelique Augustin, Christian Schori, Marc Bickle, Matthias P. Lutolf, Timothy Recaldin, Joep Beumer, Mikhail Nikolaev, Nikolche Gjorevski, J. Gray Camp

## Abstract

The intestinal mucosal barrier contains microbial organisms within the lumen while preserving the ability to absorb nutrients. Dietary, microbial, and other exposures shaped human barrier evolution and continue to impact disease susceptibility. Here, we established engineered barrier models of the human small intestine and colon composed of a multilineage epithelium, mucus layer, accessible microbial compartment and autologous tissue-resident immune cells. The epithelium has crypt- and villus-like topological domains, with stem cells differentiating into absorptive and secretory lineages with region-specific identities. Secreted mucins accumulate apically, forming a dense mucus layer separating the epithelium from colonizing commensal and pathogenic bacteria. Intestinal memory T cells integrate into and interact with the epithelium. We use the engineered intestinal tissues to identify an epithelial gene regulatory network underlying response to *Salmonella* Typhimurium infection, and uncover epithelial-immune-pathogen crosstalk coordinating cytokine release and epithelial damage. Overall, this work allows for the modular integration of epithelial, microbial, and immune compartments providing a versatile system for studying human intestinal physiology and pathologies.

## Main text

The human intestine allows the passage of nutrients and drugs, while supervising interactions with commensal and pathogenic microbes present in the lumen^1^. Controlled contact between the intestine and microbes is essential for shaping diverse aspects of human biology, including development, maturation, and specialization of both the intestinal epithelium and systemic immunity^2–5^. The intestine also provides a barrier to pathogenic species, helping contain and prevent systemic spread during infections^6^. Dysregulated intestinal-microbe interactions are a risk factor in pathologies such as inflammatory bowel disease^7,8^ and are thought to contribute to aspects of malignant transformation and cancer progression^9^.

The nuanced and context-dependent crosstalk with microbes in the intestine is executed through a multi-barrier system, comprising mature, functional and spatially patterned epithelium, an intestinal mucus layer acting as a physical and chemical barrier, and a diverse community of immune cells specialized for both tolerance and response^10^. Integration of these three components is required to meaningfully capture intestine-microbe interactions in a human-relevant setting.

Epithelial organoids derived from intestinal stem cells (ISCs) provide an inroad into functional studies of human intestinal epithelial biology^11–13^. However, intestinal organoids face limitations recapitulating intestinal physiology, owing to an inaccessible lumen, culture heterogeneity, non-homeostatic growth, lack of biomimetic morphology, incomplete cell maturation, and absence of non-epithelial intestinal lineages^14–17^. Recent bioengineering advances have overcome some of these challenges by using three-dimensional (3D) extracellular matrix (ECM)-scaffolds to provide organoids with crypt-villus epithelial architecture and access to the apical and basal side of the epithelium^18–22^. Despite these advances, physiological integration of epithelial, tissue-resident immune and microbial compartments remains a major challenge, particularly in a human context^23,24^.

Here, we establish engineered immunocompetent human intestinal models composed of autologous tissue-resident memory T cells (TRMs) and small intestinal or colonic ISC-derived epithelia that are amenable to luminal manipulation. Multiplex immunofluorescence (mIF) and single-cell RNA-seq (scRNA-seq) characterization of these barriers reveal spatially defined niches with cell type patterning along the ‘crypt-villus’ axis, reminiscent of the organ of origin. The differentiated engineered epithelium recapitulates key in vivo properties related to cell composition, barrier function, nutrient absorption, mucus layer formation, and immune-epithelium interactions. We applied the model for the study of host-pathogen interactions under *Salmonella enterica* subsp. *enterica* serovar Typhimurium (*S*.Tm) infection, providing a comprehensive single-cell multiomic view of the epithelium response to infection. Finally, we use the immunocompetent intestinal tissues to model epithelialimmune-pathogen interactions, finding that TRMs respond to *S*.Tm by cytokine release and concomitant epithelial damage, which may facilitate pathogen clearance and disease resolution.

## Results

### Biomimetic engineered human intestinal epithelium with apical access

To model human intestinal epithelium, we seeded dissociated ISC-derived organoids onto extracellular matrix (ECM) scaffolds with a surface microtopography resembling intestinal crypts (Fig. 1a-c, Extended Data Fig. 1a-b, Supplementary Table 1). Briefly, we developed a fluidic device and microstamping technology allowing us to mold an ECM-like hydrogel^25^ embedded in the core of the device with crypt-mimicking microwells (Extended Data Fig. 1a-c). The top reservoir of the device is used for cell seeding and subsequently serves as the apical medium reservoir after epithelial coverage and barrier formation. Medium from the side reservoirs passively diffuses through the gel, providing basal nutrients to the epithelial tissue. Dissociated intestinal organoids lined the hydrogel and adopted its surface topography, forming a tight epithelial barrier with regularly repeating crypts and inter-villus regions that could be maintained for at least 10 days (Fig. 1d, Extended Data Fig. 1d-f).

**Figure 1.**
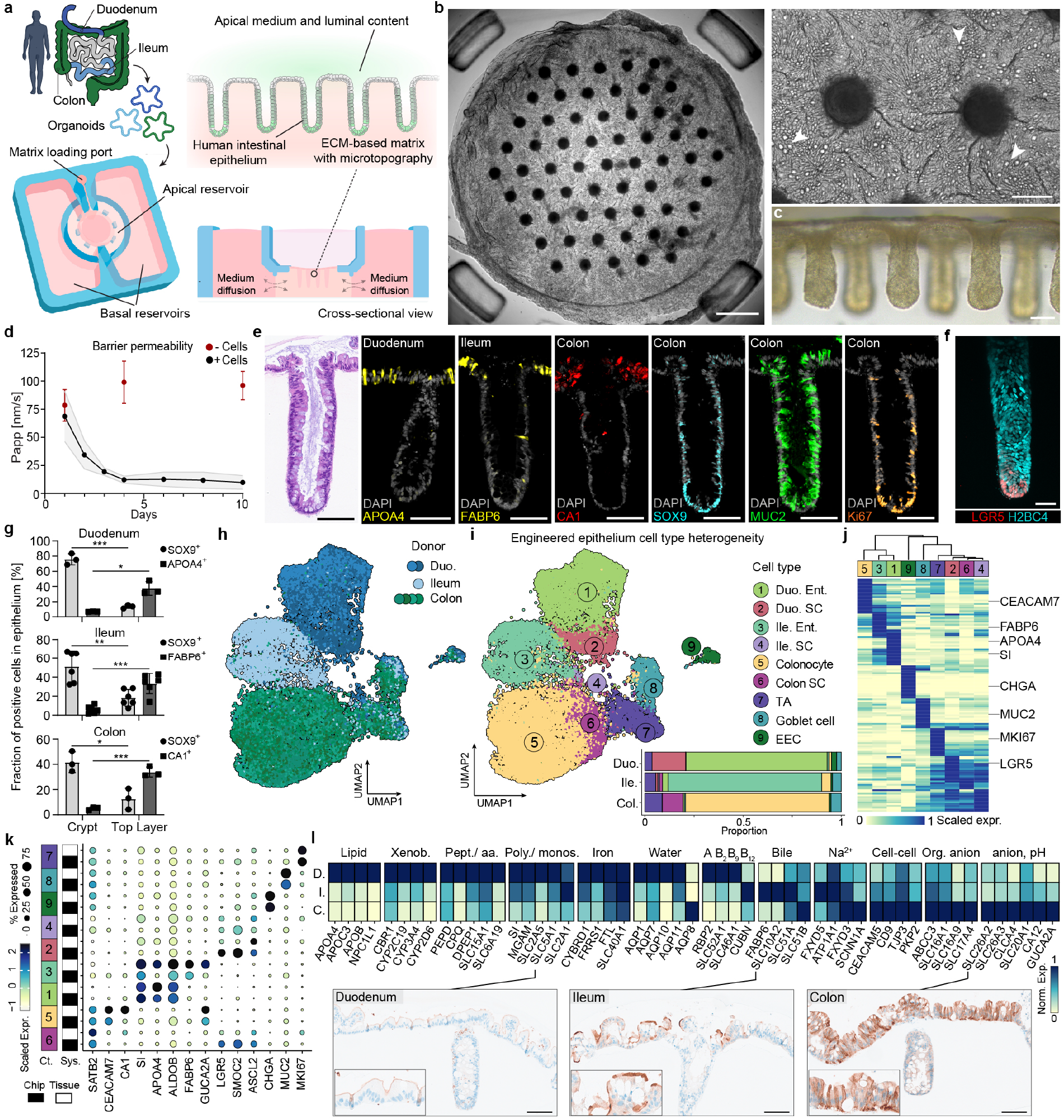
Bioengineered intestinal epithelium with biomimetic tissue topography, luminal access and region-specificity. a) Schematic overview of the engineered intestine generation from human intestinal organoids isolated from duodenum, ileum, and colon. The schematic shows the compartmentalization of the fluidic device and a cross-sectional view of the resulting intestinal epithelium. b) Representative extended depth of focus (EDF) brightfield image of a Z-stack, showing the top-view of a day-9 ileal engineered epithelium (left, scale bar: 500 µm), with crypts (large black dots) penetrating into the matrix underneath. The magnified insert on the right highlights two crypts of the epithelium, including goblet cells (arrows). Scale bar: 100 µm. c) Representative brightfield image of two rows of biomimetic crypts. Scale bar: 100 µm. d) Apparent permeability of combined duodenal, ileal, and colonic engineered epithelia over 10 days (black) compared to empty controls (red). d=1 per region, n=3-5 per donor and timepoint (+Cells), n=3-4 non-seeded chips per timepoint (-Cells). Line connects averaged P_app_ of the three donors at each timepoint. SD is represented by a grey shade. e) Representative hematoxylin and eosin (H&E) and multiplex immunofluorescence (mIF) stainings of crypt cross-sections, showing crypt-associated (SOX9+, Ki67+), secretory (MUC2+), and absorptive (enterocyte: APOA4+, FABP6+; colonocyte: CA1+) intestinal cells. Scale bar: 100 µm. f) Representative whole-mount image of iCas9 H2B4C-mTurquoise2 (cyan, nuclei) and LGR5-IRES-NLS-mScarlet3 (red) duodenal reporter-line, visualizing LRG5+ stem cells in the crypts. Scale bar: 50 µm g) Quantification of SOX9+ and absorptive lineage marker of the respective intestinal region in the crypt and top-layer of the epithelium at day 10, normalized to the DAPI+ cells within the respective annotation layer. Per intestinal region (d=1), multiple engineered intestines (n_Duo_=3, n_Ileum_=6, n_Colon_=3) were analyzed. Bar represents the mean +/-SD of the engineered intestine (dots), unpaired two-tailed t test, with Welch’s correction where indicated (***P_Duo, SOX9_= 0.0002, *P_Duo, APOA4, Welch_= 0.0279; **P_Ile, SOX9_= 0.0013, ***P_Ile, FABP6, Welch_= 0.0009; *P_Col, SOX9_= 0.0126, ***P_Col, CA1_= 0.0003). h-i) UMAP embeddings of day-10 engineered epithelia based on multiplexed scRNA-seq data. The cells are coloured by the region of origin of the respective donor (d_Duo_=2, d_Ileum_=1, d_Colon_=4; n=3 pooled per donor prior dissociation) (left) or cell type (right). The stacked barplot indicates the average proportion of each cell type in all regions. j) Heatmap showing scaled average expression of top 20 cell type-enriched marker genes of day-10 engineered epithelium from duodenum, ileum, and colon. k) Dot plot highlighting average epithelial cell-type markers from duodenum, ileum, and colon, comparing average expression between day-10 engineered epithelium and the reference primary epithelium. l) Comparison between duodenal (D.), ileal (I.), and colonic (C.) day-10 enterocyte and colonocyte gene expression for curated genes associated to the metabolism or uptake of different solutes of the intestine, normalized to the maximum expression for each gene per region (Xenob.-Xenobiotic; Pept./aa.-Peptides/aminoacid; Poly./monos.-Polysaccharide, monosaccharide; Org. anion-Organic anion; top). Representative IHC staining of SLC5A1, SLC10A2, and SLC26A2 in a single cross-section containing duodenal, ileal, and colonic day-9 engineered epithelia (combined with Extended Data Fig. 4f; bottom). Scale bar: 100 µm.

To characterize the morphology, cellular composition, and patterning of the epithelium, we developed a 3D-printed ‘histomold’ that enables simultaneous embedding of up to 50 tissues, and sectioning in a direction perpendicular to the barrier (Extended Data Fig. 1g). Hematoxylin and eosin (H&E) and multiplex immunofluorescence (mIF) staining of engineered epithelia showed histological similarity to donor-matched primary tissue in all intestinal regions, in terms of epithelial morphology and cell type patterning (Fig. 1e, Extended Data Fig. 1h, Extended Data Fig. 2a-c). In addition, we established a duodenal dual reporter line stably labeling chromatin (H2BC4-mTurquoise2) and LGR5+ stem cells (LGR5-IRES-NLS-mScarlet3), and observed LRG5+ cells localized to the crypt bottom (Fig. 1f). By day 10, a thick epithelium contained proliferative SOX9+ and Ki67+ cells primarily confined to the crypt, whereas mature enterocytes (duodenum, APOA4+; ileum, FABP6+) and colonocytes (CA1+) were enriched within the inter-crypt regions (Fig. 1g, Extended Data Fig. 3a-f). We also confirmed the presence of αDEF5+ Paneth cells in some crypts of small intestinal engineered epithelium (Extended Data Fig. 3g). These data confirm that imposing in vivo-inspired geometries on human intestinal organoids drives reproducible cell type regionalization akin to that of the intestinal crypt-villus axis^21^.

Multiplexed single-cell (sc) RNA-seq of engineered epithelia demonstrated region-specific stem, absorptive, and secretory lineages across 7 donors at day 10 of culture (Fig. 1h-j, Extended Data Fig. 4a-b, Supplementary Table 1 and 2). Cell-type composition was similar to primary counterparts and proportions were consistent across donors, highlighting the fidelity and robustness of the system (Fig. 1k, Extended Data Fig. 4c). Gene Ontology analysis of marker genes of enterocytes and colonocytes derived from the engineered duodenum, ileum, and colon tissues revealed enrichment of distinct physiological programs concordant with those of thea primary intestinal regions^26–28^ (Extended Data Fig. 4d, Supplementary Table 3 and 4). Proximal small intestinal enterocytes showed higher expression of genes involving lipid, xenobiotic, peptide, iron, polysaccharide, and vitamin metabolism or uptake (Fig. 1l; Extended Data Fig. 4d). Enterocyte maturation was confirmed by pseudotemporal ordering of the duodenal stem to enterocyte trajectory, approaching AQP10+ expression as observed at the human villus tip in vivo^29^ (Extended Data Fig. 4e). Colonocytes were enriched in cell-cell interactions and ion transport, which are critical in regulating barrier, electrolyte, and fluid homeostasis^28,30^ (Fig. 1l; Extended Data Fig. 4d). Distal small intestinal enterocytes shared some of the previously mentioned features, and showed higher expression of bile acid and vitamin B12 transport genes, reflecting their native function^31,32^ (Fig. 1l; Extended Data Fig. 4d). We confirmed differential protein expression of transporters SLC5A1, SLC10A2, and SLC26A2 between engineered epithelia of the duodenum, ileum, and colon (Fig. 1l, Extended Data Fig. 4f). Among secretory lineages, EECs and goblet cells were differentially abundant in the duodenal and colonic epithelia (Extended Data Fig. 4g-h, Extended Data Fig. 2), with substantial heterogeneity identified in EECs (CHGA+) including EC (TPH1+), MX (GHRL+, MLN+), D (SST+), K (GIP+), I (CCK+), and L (GCG+, PYY+) subpopulations that match the tissue distribution^33–35^ (Extended Data Fig. 4i-j).

Collectively, these data show that the engineered epithelium retains region-specific cell composition and gene expression. The high degree of epithelial maturation, spatial organization and regionalization, along with the basal and apical access, provide inroads to explore nutrient, xenobiotic and host-microbe processes in a relevant human context.

### Formation of a mucus layer with in vivo-like region-specific properties

The mucus, a layered network of glycosylated mucins, cellular material, antimicrobials, and other structural factors, provides an interactive barrier between the microbiome and underlying host intestinal epithelium^36,37^. To facilitate mucus accumulation into an in vivo-like layer, we reduced the apical media volume over the course of ∼4 days, which resulted in a fibrous mucus network covering the epithelium (Fig. 2a-b, Extended Data Fig. 5a-c). We also noted the presence of shed cells throughout the mucus layer, which increase mucus viscosity through DNA release^38^ and can remain viable in the gut lumen exerting an antimicrobial role^39^ (Extended Data Fig. 5c-d). Longitudinal sections along the engineered tissues revealed a mucus layer with plumes that emanate from the crypts similar to observations in primary tissue^40^ (Fig. 2c-d, Extended Data Fig. 5e). Quantification of the mucus layer thickness in these sections indicated an irregular profile, with higher thickness in the colonic epithelium (Fig. 2e, Extended Data Fig. 5f-h). Alcian Blue with Periodic Acid–Schiff (AlcianBlue/ PAS) counterstaining showed secretion and glycosylation differences between proximal and distal goblet cells and mucus, with higher ratios of negatively charged groups in colonic samples (Fig. 2f, Extended Data Fig. 5i). These patterns were comparable to those in the primary tissue and may be related to the different ratios of fucose, sialic acid, and sulfur groups across regions in vivo^41,42^ (Extended Data Fig. 5i-j). To gain deeper insight into the components of the mucus, we performed proteomic analysis of the colonic luminal content. We identified the core mucus layer proteins^43^ (MUC2, FCGBP, TFF3, AGR2, CLCA1, ZG16, and MUC5B) as major constituents, alongside additional mucins (MUC1, MUC13), antimicrobials (LCN2, DMBT1, LYPD8), and lectin-binding proteins (LGALS3, LGALS4, LGALS3BP; Fig. 2g, Extended Data Fig. 5k, Supplementary Table 5). Polymeric immunoglobulin receptor (PIGR), involved in host defense by facilitating the transcytosis of IgA and IgM immunoglobulins from the basolateral to the apical side of the epithelium^44^, was highly abundant in samples from all donors.

**Figure 2.**
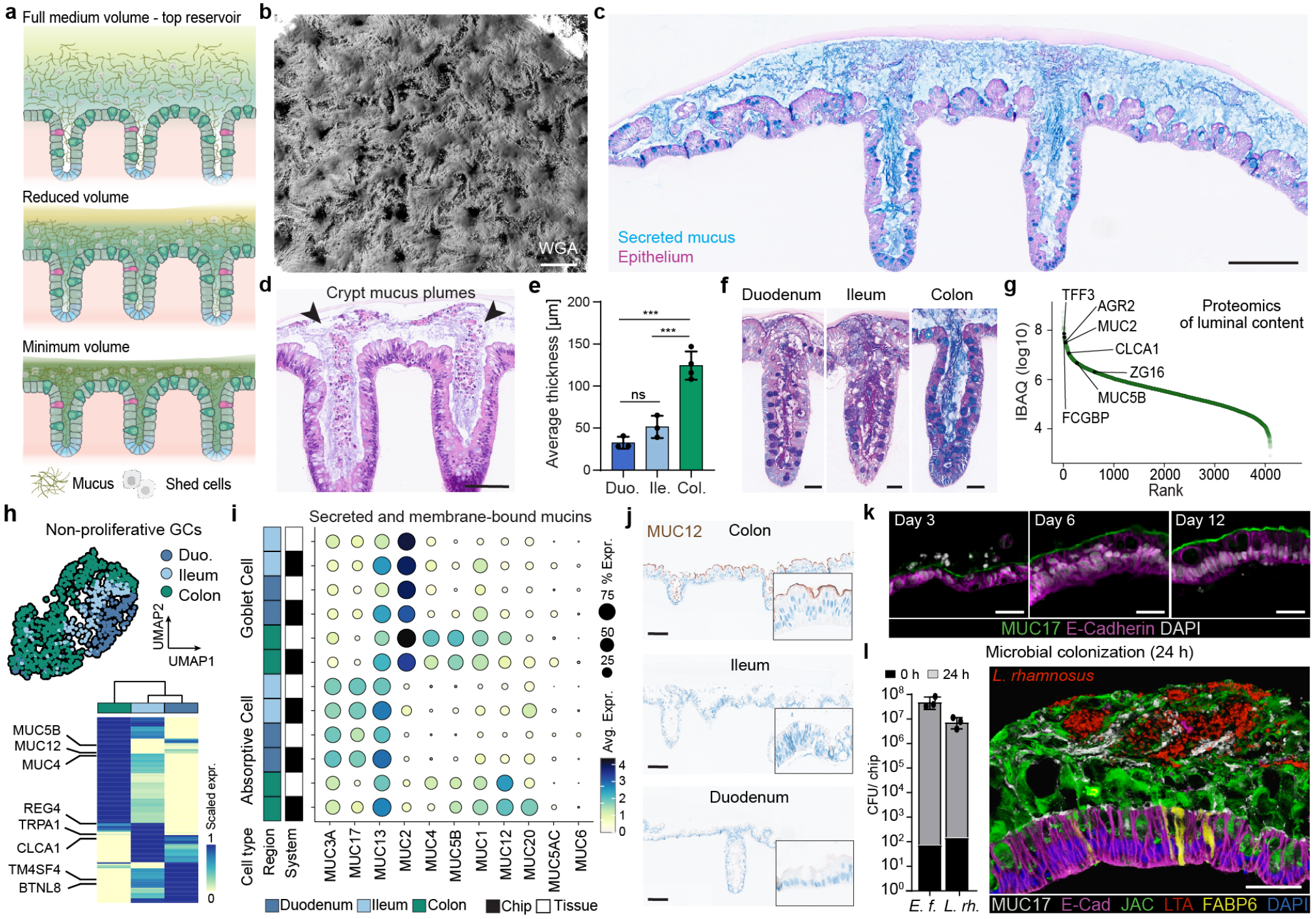
Formation of a mucus layer with in vivo-like region-specific properties. a) Schematic of the mucus accumulation strategy on the apical compartment of the engineered epithelium. Reduced apical volumes allow the accumulation of mucins and shed cells on the surface of the epithelium. b) Fluorescent live image showing a 3D reconstruction from a Z-stack of a day-13 ileal engineered epithelium with accumulated mucus (5 days, 5 µl), stained with Wheat Germ Agglutinin (WGA; grey). Scale bar: 250 µm. c-d) Representative AlcianBlue/ PAS and H&E staining of day-9 colonic epithelium cross-sections with visible goblet cells and accumulated mucus (c: 5 days, 30 µl; d: 4 days, 5 µl). Mucus plumes (arrows) appear at the crypt opening. Scale bars: 200 µm (c), 100 µm (d). e) Average thickness of the accumulated mucus layer (5 days, 30 µl) in cross-sections of day-9 multi-region engineered epithelia. Bars represent mean +/-SD of d=3-4 per region (dots) with n=3-5 averaged per donor. The mucus thickness of each engineered intestine was measured in two separate cross-sections and averaged. One-way ANOVA with Tukey’s multiple comparison test (***P_Col.-Ile._=0.0005, ***P_Col.-Duo._=0.0001, ns=P>0.05). f) Representative AlcianBlue/ PAS staining of day-9 multi-region engineered crypt cross-sections showing different glycosylation patterns between regions. Scale bar: 50 µm. g) Ranked abundances based on IBAQ values of proteins (dots) detected in the luminal content of colonic engineered epithelium with accumulated mucus (5 days, 5 µl). Core mucus layer proteins are highlighted. IBAQ values were averaged from d=3 and n=3 per donor. h) UMAP embedding of non-proliferative goblet cells (MKI67-) at day 10, with cells coloured by their region of origin (top). Heatmap showing scaled average expression of differentially expressed genes between duodenum, ileum and colon goblet cells (bottom). i) Dot plot showing average expression of the main intestinal secreted and membrane-bound mucins. Data is shown for enterocytes/ colonocytes and goblet cells of the engineered epithelium (day 10) and reference primary tissue, across regions. Data in h-i) derived from day 10 scRNA seq engineered tissue atlas (Fig. 1). j) Representative IHC staining showing the region-specific expression of MUC12 in a cross-section containing day-9 multi-region engineered epithelia. Scale bar: 100 µm. k) Representative cross-sections of duodenal epithelium, showing the development and maturation of the glycocalyx (MUC17, green) and the epithelium (ECAD, purple) over time. Scale bar: 25 µm. l) CFU counts of *Enterococcus faecium* and *Lacticaseibacillus rhamnosus* in the apical side of day-9 ileal engineered epithelium with accumulated mucus (4 days, 30 µl), colonized independently for 24 h. Bars represent mean +/-SD of d=1, n=3 (dots; left). Representative mIF staining of day-9 ileal engineered epithelium with accumulated mucus (4 days, 30 µl) after 24 h of co-culture with *Lacticaseibacillus rhamnosus* (LTA). Scale bar: 50 µm.

We utilized the scRNA-seq data to examine goblet cell (GC) heterogeneity in the engineered tissues. Among the non-proliferative GCs, we identified seven clusters (C0-C6) representing regional diversity and differentiation state (Fig. 2h, Extended Data Fig. 5l, Supplementary Table 3). Duodenum and colon GCs showed differential expression of the mucus layer components CLCA1 and MUC5B, respectively (Fig. 2h). Interestingly, ileum GCs showed expression of TRPA1, an irritant receptor with luminal sensing functions previously studied in mouse crypt EECs^45^, indicating a possible role in human GCs. We found that C1, which is dominated by colonic goblet cells, was enriched in intracellular vesicle transport, protein glycosylation and aerobic respiration, suggesting a higher mucus production machinery consistent with mucus thickness observations (Extended Data Fig. 5l-m, Supplementary Table 4). Next, we evaluated the expression of membrane-bound mucins, components of the glycocalyx that protect against bacterial attachment^36,46^. We observed a strong cell-type and region-specific expression that was comparable to that of reference tissue (Fig. 2i). For example, enterocytes in the small intestine expressed high levels of MUC3A and MUC17, whereas colonocytes expressed MUC1 and MUC12. MUC6 and MUC5AC, common stomach secreted mucins often expressed in the intestine under pathological conditions^47,48^, were nearly absent. Staining for MUC12 and MUC17 proteins in the engineered epithelium and tissue counterparts confirmed the specificity and the maturation of the glycocalyx, revealing in vivo-like expression levels by day 12 of culture (Fig. 2j-k, Extended Data Fig. 5n).

To evaluate the functionality of the mucus and the potential of the model to study host-microbe interactions, we independently colonized ileal engineered epithelia with *Enterococcus faecium* and *Lacticaseibacillus rhamnosus* – commensal members of the human gut microbiota^49,50^. After 24 hours of co-culture, both bacterial species proliferated and colonized the mucus without disrupting epithelial integrity (Fig. 2l, Extended Data Fig. 5o). Altogether, these results indicate that the engineered epithelium can accumulate a complex mucus layer with in vivo-like and region-specific properties, opening new avenues for human mucus research and the study of epithelium-mucus-bacteria interactions.

### Human epithelial response to *Salmonella* Typhimurium infection

*Salmonella enterica* subsp. *enterica* serovar Typhimurium (*S*.Tm), a common foodborne pathogen and predominant non-typhoidal *Salmonella* subtype, colonizes the gut triggering a host inflammatory response that facilitates outcompetition of the commensal microbiota^51–53^. In the lumen, *S*.Tm uses a motile flagella to reach the epithelium, where two type III secretion systems (T3SS-1 and T3SS-2) enable the invasion, survival, and expansion of the pathogen^51,54,55^. Experiments in mice have shown that *S*.Tm can penetrate the mucus layer, yet areas with discontinuous mucus are more vulnerable and prone to colonization^56^.

To study the protective role of the mucus layer against pathogenic bacteria in a human context, we infected engineered colonic epithelium with 10^5^ colony forming units (CFU) of *S*.Tm in conditions with nutrient-rich medium and different densities of accumulated mucus (Fig. 3a-c). At 18 hours post infection, *S*.Tm triggered strong epithelial damage in conditions with less mucus (Mucus^lo^), whereas the epithelium remained intact and histologically unaffected in conditions with a dense mucus layer (Mucus^hi^; Fig. 3b-d, Extended Data Fig. 6a-d). Furthermore, the dense mucus layer led to a decrease in microbial burden and preservation of the barrier integrity (Extended Data Fig. 6e-f). Nevertheless, dense mucus conditions (Mucus^hi^) showed basal secretion of IL-8 and sFAS, as well as specific zones of Caspase-3 induction that lead to increased cell shedding and accumulation of cell aggregates in the mucus layer, which appeared to impede *S*.Tm colonization of crypt locations (Extended Data Fig. 6g-h). In conditions where *S*.Tm reached the epithelium (Mucus^int^, Mucus^lo^), we could detect cells in intercrypt regions with intracellular bacteria, although the majority of the *S*.Tm load localized extracellularly, as observed in mouse models^53^ (Extended Data Fig. 6i). This inherent ability of *S*.Tm to invade and survive within epithelial cells was confirmed via live imaging using a duodenal mScarlet3-CDH1+ line and a gentamicin protection assay (Extended Data Fig. 6j-k).

**Figure 3.**
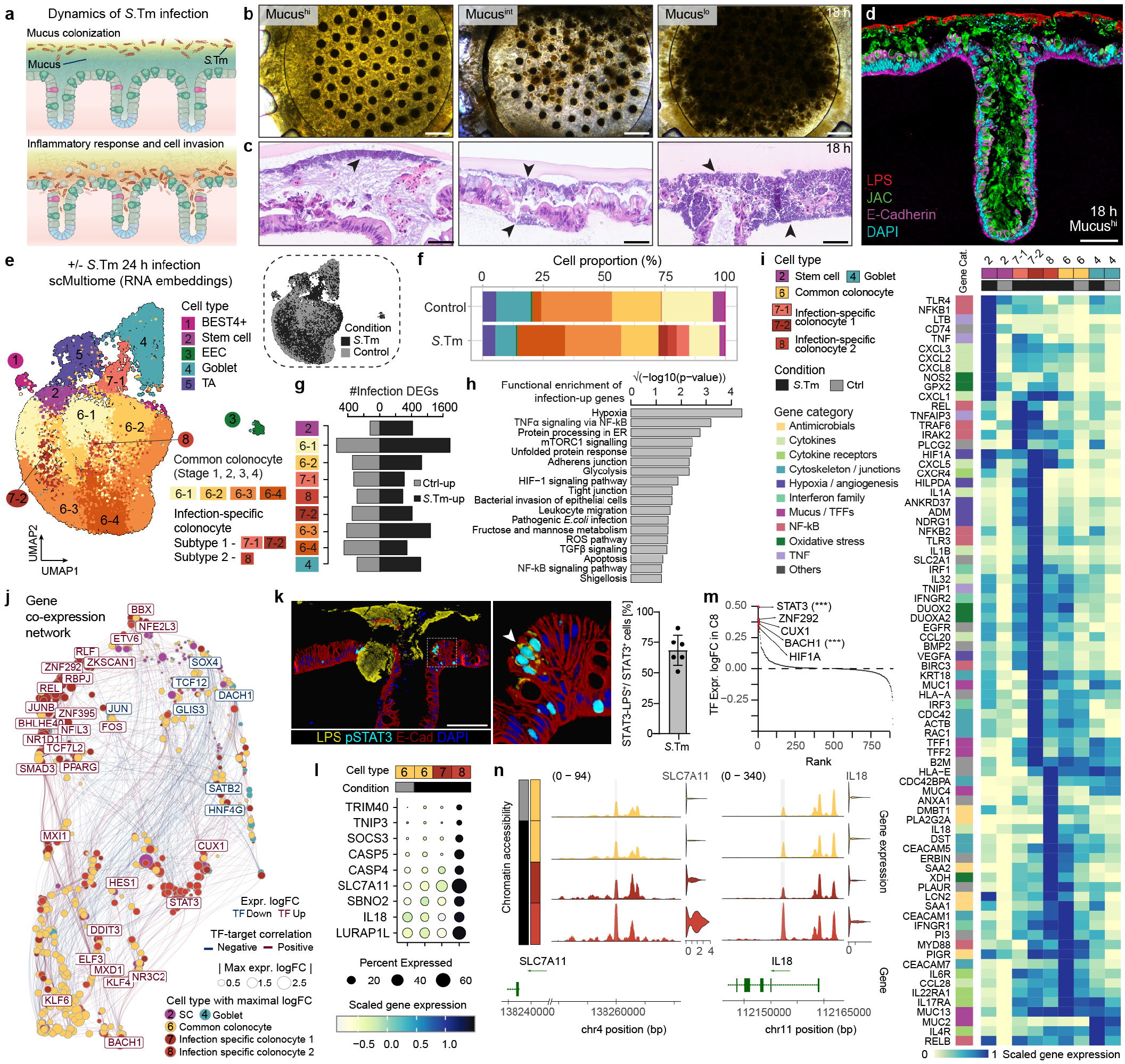
*Salmonella* Typhimurium colonization reveals cell type-specific gene regulatory response to pathogen infection. a) Schematic representation of a *S*.Tm infection and invasion of colonic engineered epithelium. b-c) Representative bright field and H&E images of day-10 colonic engineered epithelia with different levels of accumulated mucus after 18 h of *S*.Tm infection. Mucus accumulation: Mucus^hi^ (4 days, 5 µl), Mucus^int^ (2 days, 5 µl), and Mucus^lo^ (4 days, 120 µl, +wash). The degree of epithelial damage inversely correlates with the accumulated mucus level. Black arrows localize the *S*.Tm biofilm. Scale bars: 500 µm (b), 50 µm (c). d) Representative mIF image of a Mucus^hi^ day-10 colonic engineered epithelium (purple, cyan) with crypt and intercrypt mucus (green), which protects the attachment of *S*.Tm (red) to the epithelium after 18 h of co-culture. Scale bar: 100 µm. e) UMAP of single cell transcriptomic data from day-10 Mucus^hi^ colonic engineered epithelium (d=1, n=5 pooled for nuclei isolation) with and without 24 h of *S*.Tm infection, with cells colored by cell types (left) or condition (top right). By 24 h, *S*.Tm had reached and breached the epithelial barrier. f) Cell type composition of colonic engineered epithelium with and without *S*.Tm infection. g) Number of differentially expressed genes (DEGs) identified in stem cells, colonocytes and goblet cells. h) KEGG pathway and MSigDB Hallmark gene set enrichment of genes upregulated upon *S*.Tm infection. i) Heatmap showing scaled gene expression patterns of innate sensing associated genes across cell types per conditions. j) Co-expression network of top 500 DEGs and their top correlated Pando inferred transcription factors (TFs) across cell types. Edges indicate Pando predicted TF-target pairs and color of edges represent expression correlation. k) Representative mIF image and magnified insert of day-10 colonic engineered epithelium (red, blue) after 24 h of *S*.Tm (yellow) infection, showing phosphorylated STAT3 (cyan) in cells containing *S*.Tm. Scale bar: 100 µm (left). Barplot showing fraction of STAT3+/LPS+ double positive cells among the total STAT3+ cells of cross-sections from left image. Bar represents the mean +/-SD of n=6 (dots). l) Dotplot showing infection-specific colonocyte subtype 2-enriched genes across colonocyte subtypes. m) Scatter plot showing TF expression log-transformed fold change (logFC) between infection-specific colonocyte subtype 2 and matched control cells in descending order. TFs with top 5 logFC are highlighted. *** indicates significant enrichment of TF binding motifs in open chromatin regions with increased accessibility in infection-specific colonocyte subtype 2 as compared to matched control cells, BH-corrected hypergeometric test P<2.2e-06. n) Coverage plots showing normalized chromatin accessibility levels of potential distal enhancer of SLC7A11 and intronic regulatory region of IL18. Violin plots showing expression of colonocyte subtype 2 enriched genes SLC7A11 and IL18 across colonocyte subtypes in different conditions.

Next, we used single-cell transcriptome and accessible chromatin sequencing (scMultiome) to characterize the response of the human colonic epithelium with accumulated mucus (Mucus^hi^) to *S*.Tm after 24 hours of infection and epithelial breach (Extended Data Fig. 7a-b). We detected canonical cell types in control and *S*.Tm conditions with increased proportion of mature colonocytes upon infection (Fig. 3e-f, Extended Data Fig. 7c–e). The increased proportion of absorptive cells upon *S*.Tm infection was also reported in adult mouse tissue^57^. Notably, colonocyte subtype 1 (RNA cluster C7-1 and C7-2) and subtype 2 (RNA cluster C8) were nearly exclusive to the infection condition (Fig. 3f, Extended Data Fig. 7d). Overall, all cell types exhibited a pronounced response to infection, and we identified 3,056 differentially expressed genes (DEGs) and 34,337 differentially accessible open chromatin regions (DARs) between conditions (Fig. 3g, Extended Data Fig. 7f-h, Supplementary Table 3). Pathway enrichment analysis on DEGs revealed infection upregulation of hypoxia, NF-kB signalling, metabolic changes, oxidative stress, unfolded protein response, and disruption of barrier integrity (Fig. 3h, Supplementary Table 4). Infection was characterized by the upregulation of antimicrobials (LCN2, PIGR, PLA2G2A), reactive oxygen species (ROS; DUOX2, DUOXA2), angiogenic factors (VEGFA, ADM), cytokines (CCL20, IL1B, IL32), mucosal healing (TFF1, TFF2), cytoskeletal (ACTB, PLEC, DST), and membrane-associated genes (CEACAM1, CEACAM5), among others (Fig. 3i). Notably, stem cells strongly upregulated LCN2, chemokines (CXCL1, CXCL2, CXCL3), and immune-modulators (TNF, CD74, LTB). We confirmed increased protein expression of LCN2 in the epithelium, which aims to sequester iron to limit bacterial growth, yet *S*.Tm has developed mechanisms to counteract this strategy^58^ (Extended Data Fig. 7i). Altogether, these results identify a complex and heterogenous innate epithelium response involving sensing, barrier integrity maintenance, and signaling to other mucosal compartments.

Next, we used the chromatin accessibility data to infer a gene regulatory network (GRN) underlying epithelial response to *S*.Tm (Fig. 3j). This network revealed transcription factor (TF) nodes and target genes that orchestrate a broad-spectrum response, along with clusters that specifically coordinate infection-specific states. We identified BHLHE40, ZNF292, and RBPJ as coordinators of the hypoxic response observed in infection-specific colonocyte subtype 1. STAT3, integral to adaptive and innate immunity regulating pro- and anti-inflammatory responses^59^, exhibited increased gene expression in infection-specific colonocyte subtype 2. Interestingly, STAT3 has been linked to *S*.Tm infection^60–62^ and previous work has characterized the molecular mechanism by which the *S*.Tm effector SarA mediates STAT3 phosphorylation and activation^63–66^. This activation leads to an anti-inflammatory transcriptional program in the invaded cell that supports the intracellular niche and spread of the infection. Co-staining of phosphorylated-STAT3 (p-STAT3) and LPS revealed a strong co-localization, with no p-STAT3 detection in non-infected samples (Fig. 3k, Extended Data Fig. 8a-b). STAT3 activation was also detected in *S*.Tm infected cells of small intestinal engineered epithelium (Extended Data Fig. 8c). These results, in addition to the expression of the STAT3 negative feedback regulator SOCS3 in the infection-specific colonocyte subtype 2, suggest that this population contains cells with intracellular *S*.Tm (Fig. 3l).

To gain deeper insights into the responses of the different cell populations, we classified DEGs into co-expression gene modules based on expression differences across cell types between conditions (Supplementary Table 3). We found that while gene modules that mark infection-specific colonocyte subtype 1 showed the largest hypoxic and glycolysis response, the subtype 2-specific gene module exhibited increased ROS, endoplasmic reticulum stress derived from an unfolded protein response, and ferroptosis (Extended Data Fig. 8d, Supplementary Table 4). In the infection-specific colonocyte subtype 2, we observed expression of genes that suggest a possible antagonistic interaction between host cell and pathogen, with genes involved in caspase signalling and intracellular bacteria recognition (CASP4, CASP5, IL18)^67–69^, and genes with anti-inflammatory roles (SOCS3, SBNO2, TNIP3)^70–74^ (Fig. 3l, Extended Data Fig. 8e). BACH1, which regulates oxidative stress and ferroptosis^75^, was also among the top upregulated TFs in this cluster (Fig. 3m). Binding motifs of STAT3 and BACH1 were over-represented in *S*.Tm DARs (Hypergeometric test, BH-adjusted Pvalue<2.2e-06, Supplementary Table 6), indicating that both might be master regulators of the response observed in the infection-specific colonocyte subtype 2 population. Lastly, we identified chromatin regions with increased accessibility upon infection, including BACH1

TF binding sites within putative regulatory regions of IL18 and SLC7A11 in the infection-specific colonocyte subtype 2 (Fig. 3n, Extended Data Fig. 8f). Collectively, these findings identify a complex multipartite human epithelial gene regulatory response to *S*.Tm infection, confirm the STAT3 activation and subsequent regulatory role in invaded mature intestinal epithelial cells, and demonstrate how engineered intestines can be used to illuminate human responses to pathogen infection and invasion.

### Tissue-resident immune cells surveil a homeostatic intestinal epithelium

The intestine harbors the largest reservoir of immune cells in our body^76^, with highly specialized tissue-resident subsets being shaped by the local microenvironment upon extravasation^77,78^. Among these, tissue-resident memory T cells (TRMs), including intraepithelial TRMs, take permanent residence within the lamina propria and epithelium of the intestine^79^, safeguarding barrier integrity and providing rapid frontline defense against pathogens^80,81^. Previously, we introduced TRMs into ISC-derived organoids, showing that subpopulations integrate within the organoid epithelium^82^. Recognizing their intraepithelial character, we sought to build on this work by incorporating the TRMs into the engineered tissue to study reciprocal mucosal interactions.

We created apically accessible immuno-surveilled barriers by incorporating autologous TRMs into the engineered tissue (Fig. 4a-c). Briefly, we took advantage of the ECM scaffold supporting the tissues to introduce a thin layer of concentrated TRMs derived from proximal intestinal resections (Extended Data Fig. 9a-b). Owing to their tissue-resident phenotype facilitating epithelial interaction (Fig. 4e, Extended Data Fig. 9c-d), the majority of TRMs infiltrated the engineered epithelium at rates consistent with those of donor-matched parental samples (Fig. 4f-i, Extended Data Fig. 9e), without compromising the formation or integrity of the epithelial barrier (Extended Data Fig. 9f). In contrast, blood-derived T cells barely integrated into the epithelium, even after 9 days of co-culture (Fig. 4f). In addition to intraepithelial integration, TRMs actively patrolled the barrier at speeds similar to those previously reported^82–84^ (mean = 5.7 µm/min; Fig. 4d, Supplementary Video 1-3). Interestingly, we detected slower (mean = 3.7 µm/min) yet more persistent TRM movement within the crypt structures compared with the inter-crypt spaces (Fig. 4d, Supplementary Video 1-2).

**Figure 4.**
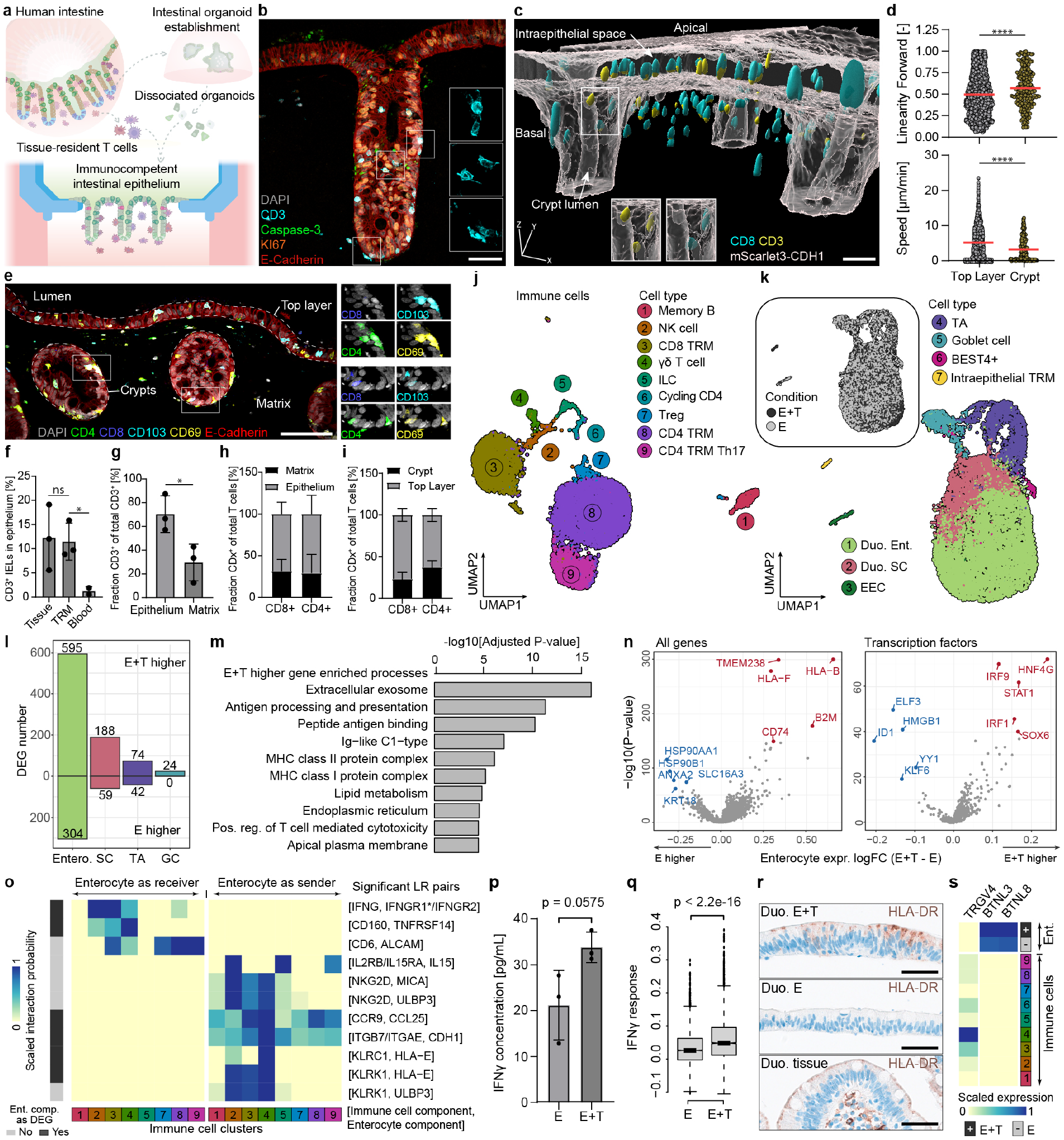
Engineered human intestinal mucosa with autologous tissue-resident T cells. a) Schematic of tissue-resident immune cell incorporation into the basal compartment of the engineered intestine. b) Representative mIF image of a crypt cross-section showing CD3+ TRMs (cyan) residing in the matrix and homeostatically (Caspase-3-, green) integrating into the epithelium (E-Cadherin+, red) at day 9. Scale bar: 50 µm. c) 3D reconstruction of CD8+ (cyan) and CD3+ (yellow) TRMs residing in the matrix and Cas9 CDH1-mScarlet3 reporter-line intestinal epithelium (E-Cadherin, grey) at day 7 (scale bar: 50 µm). d) Comparison of TRM movement speed and linearity between the top-layer and the crypt from imaging time-lapse (42 min) of the immunocompetent co-cultures. Quantification based on the respective regions of engineered intestines (n=3) derived from a single donor in a representative experiment. Dots represent the sum of single T cells detected in all engineered epithelia, the red line represents the mean. Unpaired two-tailed Mann-Whitney test (****P<0.0001).e) Representative mIF image CD4+ (green) and CD8+ (blue) expressing tissue-resident markers (CD69+, yellow; CD103+, cyan) within the matrix and intercalated in the epithelium (E-Cadherin+, red). Scale bar: 100 µm. f) Comparison between the intraepithelial integration of autologous tissue-derived, allogeneic blood-derived T cells in engineered tissues and non-matched primary tissue. Unpaired two-tailed t test with Welch’s correction where indicated (*P_Welch_=0.0359, ns=0.8749). g) Quantification of tissue-resident CD3+ T lymphocyte distribution between epithelium and matrix, unpaired two-tailed t test (*P=0.0319). h) Representative mIF image and quantification of localization (matrix vs. epithelium) and i) integration rate of tissue-resident CD4+ and CD8+ T cells in the top-layer and crypts of the engineered epithelium. Bars in f-i) represent the mean of donors (d=3) +/-SD at day 9, derived from three engineered intestines (n=3) per donor. j) UMAP embedding of scRNA-seq transcriptome data on TRMs harvested from immunocompetent engineered tissues at day 7 of co-culture, colored by cell cluster (d=1, n=6 pooled). k) UMAP embedding of scRNA-seq transcriptome data on epithelium harvested from identical immunocompetent engineered tissues and epithelium-only controls after seven days of, colored by cell cluster (d=1, n=6 pooled). l) Number of differentially expressed genes detected in the four major epithelial cell types. m) DAVID^131,132^ reported functional enrichment of genes upregulated in enterocytes upon TRM co-culture. n) Volcano plot showing differential expression of all genes and transcription factors (TF) in the enterocytes upon TRM coculture. Labels in red and blue colors highlight the up- and down-regulated genes with the highest Wilcoxon rank sum test significance respectively. o) Heatmap highlighting scaled CellChat-derived interaction probability of directed mucosal ligand-receptor pairs between enterocytes and TRMs. p) IFNγ secretion detected in the supernatant of the immunocompetent engineered tissues (day 8). Bars represent the mean +/-SD of three donors (d=3, dots), with independent engineered intestines (n=3) averaged per donor, two-tailed unpaired t test with Welch’s correction. q) Seurat derived module score based on MSigDB IFNγ response signature score between immunocompetent barrier and epithelium-only culture. r) Representative IHC staining of HLA-DR (brown) on duodenal engineered epithelia (day 9; E+T, E) and primary tissue. Scale bar: 50 µm. s) Heatmap showing expression patterns of BTNLs in enterocytes and receptor TRGV4 across immune cell types.

Next, we used scRNA-seq to profile the TRM populations within the engineered tissue, and examine TRM effects on the epithelium after one week of co-culture. TRMs preserved striking heterogeneity after one week of co-culture, comprising populations of CD8 T cells (CD8 TRMs), helper CD4 T cells (CD4 TRMs), regulatory T cells (Treg), T helper 17 cells (Th17), γδ T cells, natural killer (NK) cells, innate lymphoid cells (ILCs) and memory B cells (Fig. 4j, Extended Data Fig. 9g). We note that some of these populations (γδ T cells, ILCs, B cells) are not tissue-resident memory T cells. However, owing to the consistently highest abundance of TRM cells in the isolates across donors^82^, we use the ‘TRM’ abbreviation throughout this manuscript to refer to the pool of immune cells collected from intestinal tissue resections. In line with their long-lived, quiescent states within native tissue^85,86^, TRMs appeared to be predominantly non-proliferative, aside from a small population of cycling cells (Fig. 4j, cluster 6). While overall cell type composition and canonical epithelial markers were not affected (Fig. 4k, Extended Data Fig. 9h-j), a total 1,014 differentially expressed genes (DEGs) in response to TRM presence were identified in diverse epithelial cell types (Fig. 4l, Extended Data Fig. 9k, Supplementary Table 3). Gene Ontology enrichment analysis on differentially expressed genes (DEGs) of enterocytes revealed the induction of transcriptomic programs involving antigen processing and presentation (Fig. 4m, Supplementary Table 4), with the upregulation of MHC-I molecules (HLA-F, HLA-B, B2M) and CD74 (Fig. 4n), critical for the assembly and trafficking of MHC-II complexes^87^. TF expression analysis in the enterocyte cluster linked gene expression changes to the activation of STAT1, IRF1 and IRF9, likely induced by the interferon (IFN) signaling pathway^88–90^ (Fig. 4n). Given that IFNs, particularly IFNγ, are key orchestrators of intestinal immunity^91–93^, we hypothesized that this immune-readiness signature might be facilitated by TRM-secreted IFNγ. Indeed, ligand-receptor pairing analysis using the scRNA-seq data identified enterocytes as receivers of IFNγ from multiple TRM subsets, primarily NK and γδ T T cells as well as CD8 TRMs (Fig. 4o, Extended Data Fig. 9l). We confirmed the presence of IFNγ within the culture supernatants at the protein level (Fig. 4p), reflected by a global IFNγ-response in the immunocompetent epithelium (Fig. 4q). We also detected higher protein levels of the IFNγ signaling target HLA-DR within TRM-containing barriers (Fig. 4r). While these TFs are often associated with disease^94–97^, we did not detect a stress-related gene signature in immunocompetent epithelium (Extended Data Fig. 9m), suggesting that IFNγ drives immune competence of the system, without triggering an active immune response at this stage of co-culture.

We further confirmed canonical epithelial-immune interaction pairs within the engineered tissues, including CDH1-ITGAE98 (E-Cadherin-CD103), CD6-ALCAM99,100 (CD6-CD166) and TNFRSF14-CD160101,102 (HVEM-CD160; Fig. 4o, Extended Fig. 9l, Supplementary Table 7). In addition, we found evidence for mutually beneficial, region-specific and self-reinforcing immune-epithelial crosstalk. For example, epithelial expression of IL-15 and trans-presentation via IL-15Rα as well as CCL25 suggests that the duodenal intestinal barrier sustains and recruits CCR9+ TRMs103–107 (Fig. 4o, Extended Data Fig. 9n), whereas colonic barriers secrete the highest level of lymphocyte chemoattractant CCL28108 (Extended Data Fig. 9n). Furthermore, TRMs appear to significantly drive the upregulation of butyrophilin-like (BTNL) molecules BTNL3 and BTNL8 within the epithelium (Fig. 4s), potentially in a HNF4G-dependent manner^109^ (Fig. 4n). Epithelial BTNLs signal back to the immune compartment, driving the maturation of Vγ4+ γδT cells, which, in turn, have various protective roles in the intestine^110^. Indeed, the loss of BTNLs and HNF4G is associated with intestinal pathologies^111,112^.

Together, these data suggest that engineered intestinal tissues capture a homeostatic immune state over extended culture, preserving physiological immune cell behaviors and diversity, and fostering in vivo-like epithelial-immune interactions.

### Immune-epithelial crosstalk coordinate a response to infection

Studying human intestinal immune response to bacterial pathogens in mechanistic detail is challenging, because individuals suffering infections rarely undergo endoscopic examination and biopsy collection. IELs and TRMs are described as rapid responders to pathogenic intestinal infection in mice^83,113–116^. We took advantage of the modularity and experimental complexity afforded by the immunocompetent intestinal epithelium to study the TRM response to an acute *S*.Tm infection in a human-relevant context.

To this end, we extended the experimental infection period by lowering the initial inoculum (around 10^2^ CFU) and using a reduced medium apically (Extended Data Fig. 10a). Additionally, we controlled the invasion of *S*.Tm into the basal reservoir by supplementing the basal medium with gentamicin. Thus, we infected immunocompetent and epithelium-only duodenal barriers for 48 hours and verified the presence of *S*.Tm and TRMs on the apical and basal sides of the epithelium, respectively (Fig. 5a-b, Extended Data Fig. 10b). TRM presence did not affect bacterial growth kinetics within the system (Extended Data Fig. 10c). Cross-sectional views of infected samples after 48 hours revealed *S*.Tm biofilms that attached to the apical top-layer of the epithelium and induced epithelial damage (Fig. 5c, Extended Data Fig. 10d). We observed large numbers of intraepithelial TRMs in close proximity to the invading bacteria, in some cases only 3 µm apart (Fig. 5d, Extended Data Fig. 10e). Notably, TRMs appeared to accelerate and exacerbate epithelial damage upon infection, as evidenced by an increase in epithelial cell shedding, epithelial thinning, and permeability in immunocompetent barriers compared with epithelium alone (Fig. 5e, Extended Data Fig. 10e-f). As the infected engineered epithelium upregulates genes encoding for, among others, chemokines and cytokines specific to T cell recruitment (CXCL10, CCL20, CCL28) and effector function (IL-18, IL-1B; Fig. 3i, Extended Data Fig. 6g), we aimed to determine whether the elevated damage in the immunocompetent system could be attributed to TRM activation. We assessed the presence of T cell-related cytokines within the supernatant and observed a strong induction of IFNγ (Fig. 5f, Extended Data Fig. 10g), which orchestrates both adaptive and innate responses via diverse mechanisms^91,117^, but has also been shown to induce apoptosis of intestinal stem cells^118,119^. *S*.Tm infection also led to the upregulation of GzmB (Fig. 5g, Extended Data Fig. 10h), a hallmark of cytotoxic activation of T cells and a known mediator of immune-mediated intestinal epithelial cell shedding^120^. Further, we observed the increased production of IL-17A and IL-6 (Fig. 5h-i, Extended Data Fig. 10i-j), which, in addition to mediating various immune processes^121,122^, have roles in microbial response and resolution by inducing anti-microbial peptides within the epithelium and promoting epithelial regeneration^123–127^. We note that the upregulation of IL-6 and IL-17A was donor-dependent, potentially reflecting physiological heterogeneity within TRM populations isolated from different donors^82^. Utilizing our scMultiome dataset on *S*.Tm-infected colon epithelium (Fig. 3), we found genes encoding for cognate receptors for these cytokines (IFNγR1, IFNγR2, IL17AR, IL-6R) upregulated within the epithelium in response to the pathogen (Fig. 5j).

**Figure 5.**
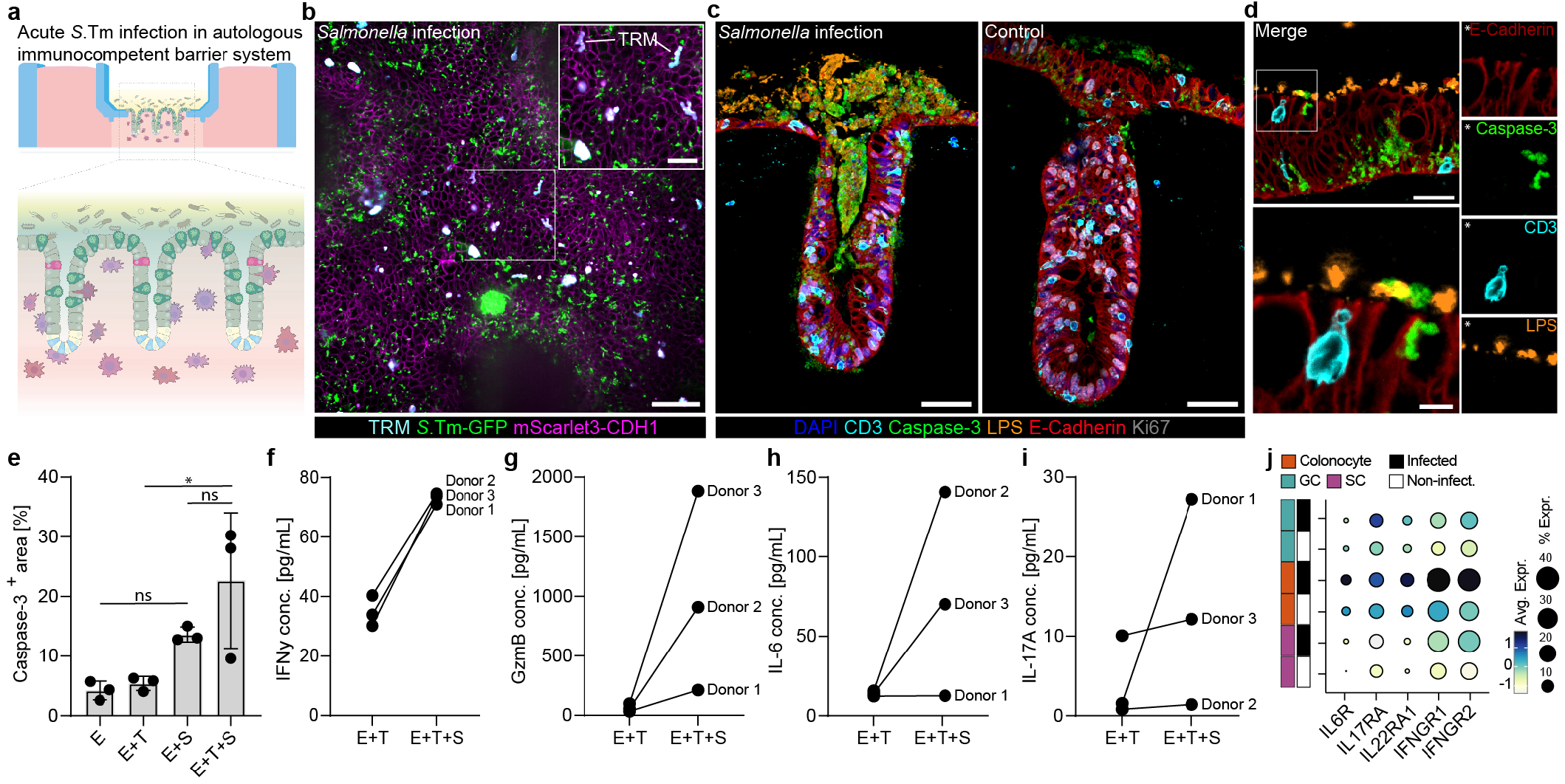
*Salmonella* Typhimurium triggers tissue-resident lymphocyte activation in immunocompetent engineered human gut tissue. a) Schematic overview of an tri-compartmental culture of a luminal pathogen, engineered epithelium and autologous tissue-resident T cells (TRMs) b) Representative EDF fluorescent image of selected Z-stacks with *S*.Tm (green) in the lumen of the Cas9 CDH1-mScarlet3 reporter-line (E-Cadherin, magenta) with intraepithelial TRMs stained with Cell Trace Far Red (cyan) at day 9. Scale bar: 50 µm (main), 25 µm (insert). c) Representative mIF images exemplify LPS+ *S*.Tm-mediated (orange) apoptosis (cleaved Caspase-3+, green) in the duodenal engineered epithelium (E-Cadherin+, red; Ki67+, grey) with intraepithelial CD3+ T cells (cyan) at site of infection after 48h (day 9). Scale bar: 50 µm. d) Magnified insert of a representative mIF image highlights a single CD3+ T cell (cyan) at the tip of the epithelial brush border (E-Cadherin+, red), several micrometers apart from LPS+ *S*.Tm (orange). Epithelial apoptosis detected in the basolateral side of the epithelium (cleaved Caspase-3, green). Scale bar: 20 µm, magnified insert: 5 µm. e) Quantification of apoptosis in the immunocompetent intestinal epithelium 48 h after the start of the infection (day 9). Barplot represents the mean +/-SD of the donors (d=3), two-way ANOVA with Tukey’s multiple comparisons, non-significant (ns = P>0.05) unless otherwise specified (*P=0.0383). f-i) Cytokines detected in the basal supernatant of the intestinal barrier after 48 h of infection (day 9). Each dot indicates the averaged engineered intestines (n=3-5) per donor (d=3) between infected and non-infected conditions. j) Dot plot showing average gene expression of cytokine receptors (matching cytokines secreted in f-i) in infected and non-infected colonic engineered epithelium after 24 h of *S*.Tm co-culture. Data displayed is derived from scMultiome dataset of Fig. 3.

Overall, these data suggest that pathogenic infection of human engineered intestinal tissues drives concurrent and mutually reinforcing responses within the epithelial and immune compartments, whereby intestinal immune cells are recruited, produce pro-inflammatory and damage-associated cytokines, and epithelial cells potentiate the effects by upregulating cognate receptors.

## Discussion

We report novel bioengineered intestinal tissues that comprise a luminally-accessible epithelial barrier of multiple intestinal regions, a stable layer of mucus, and autologous tissue-resident immune cells (TRMs). The engineered epithelia feature periodically repeating crypt units that maintain LGR5+ stem cells, in vivo-like cell composition and patterning, and retain transcriptomic signatures consistent with the intestinal region of origin across donors. Importantly, identical medium and physical culture conditions result in high fidelity region-specific signatures due to stem cell origin and biophysical attributes of the system. Maturation of the absorptive and secretory lineages and a reduction in total apical volume enable the formation of a mucus layer that supports commensal bacteria co-culture and features region-specific properties comparable to those of human tissue. Additionally, TRMs introduced into the engineered tissues undergo spontaneous intraepithelial integration, patrol the barrier, and engage in multi-layered and bidirectional crosstalk with the epithelium, conferring an immune responsive state within epithelial cells via IFNγ signaling. Some of these interactions have not been described before. For example, we show that TRMs themselves drive the induction of BTNLs within the epithelium, which may be self-serving, given the role of BTNLs in the selection and maintenance of the Vγ4+ γδ T cell population^128^. The engineered intestinal tissues could be used to elucidate the mechanisms underlying the TRM-mediated BTNL upregulation, which are not understood.

The modularity of the system, combined with multi-omic analyses, allowed new inquiry into *S*.Tm infection in a human context. We demonstrate that the mucus layer thickness associates with protection from pathogenic breach, subsequent damage, and invasion of the epithelium, consistent with in vivo models^56^. The recapitulation of this complex infection phenomena enabled us to map the transcriptomic signature and gene regulatory mechanisms that underlie epithelial responses to *S*.Tm at a single-cell level. We learned that the epithelium orchestrates cell type-specific and multifaceted inflammatory, metabolic, and structural responses that serve *S*.Tm^52,53^ and demonstrate the epithelium’s inherent innate-sensing capabilities. STAT3 was identified as a major gene regulatory hub, and its confirmed phosphorylation in invaded mature human epithelial cells supports previous work with immune cells and immortalized lines^60–66^. Additionally, we uncovered an interesting set of genes and TFs previously not directly linked to *S*.Tm infection that may influence the pathogenesis and intracellular niche. Investigating the functional role of these genes, including the downstream effects of STAT3 activation, would be interesting subjects for future studies.

We also identify a clear functional effect of TRMs upon *S*.Tm infection, finding that they accelerate epithelial damage and shedding, presumably via the induction of pro-inflammatory cytokines and cytotoxic granzymes. Indeed, this could be a part of the defense response and facilitate pathogen clearance, as intraepithelial lymphocytes have been shown to guard the epithelium by promoting shedding via granzyme-B-mediated ejection of stressed and apoptotic epithelial cells^129^. Importantly, our study suggests that lymphocytes, which are largely associated with antigen-specific adaptive responses, can mount innate-like responses to infection. Exploring the population-specific mechanisms that underpin these responses in the absence of professional antigen presenting cells would be interesting.

Even though this study utilized *S*.Tm to study host responses to infection, the engineered intestinal tissues and analytical framework described can be used to dissect epithelial and immune responses to other pathogens or commensal microbes of the intestine. Indeed, the model and its components expand what can be addressed ex vivo beyond questions related to intestine-microbe interactions. The mature, patterned and regionalized epithelium unlocks research directions in region-specific absorption and metabolism of drugs and nutrients, as well as mucus biology. The immunocompetent intestinal barriers could help provide previously elusive insights into the interaction between the epithelium and intraepithelial lymphocytes, which have been implicated as crucial safeguards of intestinal homeostasis, and, upon dysregulation, drivers of inflammatory bowel disease^110,111,130^. Altogether, the engineered intestinal tissues described here bring the field steps closer to capturing complex, organ-level processes in a human setting that is controllable, observable and tractable.

## ACKNOWLEDGEMENTS

We thank Alex Zipperer for the initial support with bacterial work and laboratory space; Miguel Camacho Rufino for artwork support; Barbora Lavickova for experiment discussions and results interpretation; Anke Gehringer for logistic support. We thank Regine Gerard for help with organoid and TRM isolation and establishment. We thank the Pathology, Genomics and Proteomics Core Labs of pRED for their support with infrastructure, sequencing and proteomic experiments

## AUTHOR CONTRIBUTIONS

R.L.S., M.F.H., Q.Y., M.N., N.G. and J.G.C. conceived the study; R.L.S., M.F.H., Q.Y., N.G., J.G.C. wrote the manuscript; M.N. designed and fabricated the fluidic device and crypt stamps, with support from J.A.; M.P.L. contributed to the development of the fluidic device holding the tissues; M.N. generated artwork design; M.F.H. and R.L.S. designed and performed most experiments in the manuscript; R.L.S., M.F.H., M.N., N.A. and J.B. optimized epithelium growth conditions; M.F.H. isolated organoid lines and TRMs from primary tissue; R.L.S., M.F.H., M.A.B. and L.G.T. performed scRNAseq and scMultiome experiments; M.A.B. and L.G.T. generated sequencing libraries; Q.Y. performed the computational analysis of scRNAseq and scMultiome data, with support from R.L.S., and data interpretation with R.L.S and M.F.H.; M.F.H., R.L.S., and J.A. co-developed the histomold; I.C. performed FFPE processing, sectioning, H&E and AlcianBlue/PAS stainings, with support from M.F.H.; M.F.H. performed immunohistochemistry and mIF stainings, imaging and analysis with support from R.L.S. and M.N.; M.F.H., R.L.S., M.N. and L.K. performed whole mount stainings and imaging; K.K. performed flow cytometry experiments and analysis; A.M.F. performed cytokine measurements; N.A., Y.B. and J.B. generated reporter lines; S.M. performed rheology measurements with support of M.N.; J.F., A.A. and C.S. performed and processed proteomic measurements; Live-imaging was performed by M.F.H., R.L.S. and M.N., and analyzed by M.B.; Liveimaging videos were generated by M.N.; T.R. supported with data interpretation; All authors read and approved the manuscript.

## DATA AVAILABILITY

Raw and processed single cell data with cell type annotation, statistically significant differentially accessible chromatin regions upon *S*.Tm infection and TF-target regulomes inferred by Pando based on the *S*.Tm infection dataset are available upon request.

## CODE AVAILABILITY

The core scripts of single cell data analysis are available at https://github.com/devsystemslab/Host-microbe_interaction_model/.

## Methods

### Human samples and Ethics statement

Human intestinal tissue resections, along with concurrent data collection and experimental procedures, were conducted within the framework of the non-profit foundation HTCR (Munich, Germany) and University Center for Gastrointestinal and Liver Disease (Clarunis; Basel, Switzerland), including informed patient consent. The HTCR Foundation’s framework received approval from the ethics commission of the Faculty of Medicine in the Ludwig Maximilian University (no. 025-12) and the Bavarian State Medical Association (no. 11142). The framework of the University Center for Gastrointestinal and Liver Disease was approved in accordance with the Helsinki Declaration and reviewed and approved by the ethics committee (Ethics Committee of Basel, EKBB, no. 2019-02118). Twenty consenting patients underwent visceral surgery with partial resection of intestine (precise intestinal regions are listed in Supplementary Table 1) for various oncologic indications. We used micro- and/or macroscopically tumour-free regions of resectates for further preparation and downstream handling. 16-week developing human tissue was obtained through Advanced Bioscience Resources (ABR). Consent was given by the mother and tissue was anonymized. All experiments were performed following relevant guidelines and regulations.

### Isolation of tissue-resident immune cells and intestinal organoids

TRM and crypt organoid generation was performed as previously described^82^. Briefly, intestinal tissue was processed by removal of the underlying muscularis, serosa and fat from the basal side of the intestinal tissue using forceps and scissors. The remaining mucosal tissue was thoroughly washed with PBS (10010-015, Gibco; supplemented with penicillin-streptomycin (1500 U/mL; 15140122, Gibco), gentamicin (500 µg/mL; 15750060, Thermo Fischer Scientific) and Primocin (50 µg/mL; ant-pm1, Invivogen) before using a glass coverslip to remove excess mucus from the luminal side and blood vessels from the basal side. The cleaned tissue was then cut into square pieces (∼5 x 5 mm3) with a scalpel. Next, each fragment was loaded onto a 9 x 9 x 1.5 mm3 tantalum-coated, carbon-based scaffold (MSPP23281-1-REV-A, Ultramet) and cultured in 24-well plates (CC7682-7524, CytoOne) containing 2 mL of cytokine-free immune-media (Supplementary Table 1). Twenty-four hours later, scaffolds with tissue on were removed and egressed cells were harvested from the bottom of the wells by thorough pipetting. Cells were then counted and biobanked in CryoStor (C2874, Sigma-Aldrich) before further use.

For isolation of donor-matched crypts for organoid establishment, an adjacent clean tissue piece (∼3 cm2) was used. Intestinal villi were removed from the tissue by using a glass coverslip. The tissue was then incubated on ice in PBS + 10 mM EDTA (15575020, Thermo Fischer Scientific) on a shaker for 30 min. Next, a coverslip was used to scrape off the crypts of the tissue. Crypts were collected in organoid-media (Supplementary Table 1). Crypts were centrifuged and resuspended in GFR Matrigel (356231, Corning). After polymerization for 15 min at 37°C, organoid-media see Supplementary Table 1) was added with 10 µm Y-27632 (72305, Stemcell Technologies). Following organoid formation, organoids were passaged every 7 days (up to a maximum of 25 passages). After three passages, organoid mycoplasma tests (LT07-710, Lonza) were performed. All lines used in this study were verified as negative for mycoplasma. Organoid banking was performed using CELLBANKER 1 (11888, Nippon Zenyaku Kogyo., LTD).

Passaging was performed as previously described^133^. Briefly, organoid-media (Supplementary Table 1) was removed and wells were washed with DPBS. Next, 500 µl of Gentle Cell Dissociation reagent (07174, StemCell) was added and incubated for 2 min at RT before collecting the domes in a Falcon tube. After further 15 min of incubation at RT, the suspension was centrifuged at 350 g at 4°C for 5 min. After removal of liquid, the pellet was harshly resuspended with organoid-wash-media (Supplementary Table 1) to break the organoids. Centrifugation was repeated (at 500 g) and the supernatant discarded (if needed, organoid dissociation was repeated), before resuspending the dry pellet with GFR Matrigel.

### Flow cytometry of isolated immune cells

After isolating immune cells from the tissue, the harvested cells were filtered through a 40 µm filter (43-50040-51, pluriselect) and phenotyped by flow cytometry prior to cell banking. The cells were transferred into a V-bottom plate and centrifuged at 300 g for 4 min. Afterwards, excess media was removed and 30 µL live-dead stain (65-0865-14, Thermo Fischer) was applied, followed by 10 min incubation at RT. Next, 120 µL of FACS buffer was added to each well and the centrifugation repeated. Subsequently, 50 µL of the primary antibody cocktail was applied and incubated for 15 min at RT (Supplementary Table 1). The centrifugation step was repeated before fixation with 4% PFA for 25 min at RT. The stained cell suspensions were then acquired on a Cytek Aurora with the 5L configuration (Cytek) and analysed using FlowJo v.10.

### Generation of reporter lines

Targeting vectors for LGR5-IRES-NLS-mScarlet3, CDH1-mScarlet3 and H2BC4-mTurquoise2 for homology-directed repair (HDR) were designed as described previously^134^. Human intestinal organoids were incubated with 1.25% DMSO %V/V and 10 µM Y-27632 (72305, Stemcell Technologies) in culture media 24 h prior to electroporation. Organoids were harvested, dissociated using TrypLE (12604-013, Gibco) to clumps of approximately 5 cells. 106 cells were resuspended in High Performance Electroporation Solution (BTXpress) containing up to 20 µg DNA at a 1:3 molar ratio of pCMV-SpCas9-U6-gRNA expression vector and targeting vector. Cells were transfected using the NEPA21 Super Electroporator (Nepagene) with settings as described before135. 10-14 days post transfection, successfully modified organoids were manually picked based on fluorescence signal or by sorting reporter-positive cells using fluorescence activated cell sorting (FACS). Site-specific integrations were confirmed by Sanger sequencing of PCR-amplified genomic DNA. Clones with site-specific integrations were pooled to generate polyclonal lines.

### Microdevice design and fabrication

During the work on this project, a large number of microdevice prototype versions were fabricated using a wide range of microfabrication techniques, including cleanroom soft lithography, 3D printing, two-photon polymerization (2PP) lithography, CNC milling and plastic injection molding.

The final chip is designed in a slide format (75 x 25 mm) containing three independent chips per slide. Each chip contains a central hydrogel chamber flanked by two side media reservoirs and one top media reservoir. The hydrogel precursor is injected through a loading port leading to the central gel chamber. A cylindrical stamp with a microtopography (e.g., an array of biomimetic micropillars) on its base is placed onto a stamp-holding edge of the top reservoir before gel injection. After gel polymerization (see details in a separate section), the stamp is removed, leaving an inverse microtopography (e.g., an array of microwells) on the hydrogel surface. The top and side reservoirs are then filled with PBS or cell culture medium. The top reservoir is used for cell seeding and subsequently serves as the apical medium reservoir after epithelial coverage of the hydrogel and barrier formation. Medium from the side reservoirs passively diffuses through the gel, supplying the epithelial tissue with nutrients from the basal side.

Early prototypes were fabricated using conventional soft lithography and PDMS molding workflows established at the Center of Micronanotechnology (CMi, EPFL). The 3D chip design was split into two photomasks that were sequentially aligned to enable the creation of pseudo-3D features and different heights for phase-guiding micro-pillars. Chip layouts were patterned onto chrome-coated fused silica substrates via laser lithography (VPG200, Heidelberg Instruments). Three layers of SU8 GM1075 (Gerlteltec) were sequentially spin-coated onto dehydrated silicon wafers to achieve a total mold thickness of 750µm. The first two 250-µm layers were soft-baked, exposed to UV through the first photomask (MA6/BA6, Süss MicroTec), and post-baked. Then the third SU8 layer was spin-coated, soft-baked and the second photomask was aligned to the existing pattern using dedicated alignment marks. After a final post-exposure bake and development with PGMEA (Propylene Glycol Monomethyl Ether Acetate, Sigma), the molds were hard-baked at 135°C for 4h and silanized overnight using vapor-phase trichloro(1H,1H,2H,2H-perfluorooctyl) silane (Sigma-Aldrich). PDMS (Sylgard 184, Dow Corning) was mixed at a 10:1 base-to-curing agent ratio, poured onto the molds, degassed under vacuum, and cured at 80°C for 24h. PDMS replicas were cut, punched, and plasma-bonded to glass-bottom dishes.

The strength of the soft lithography pipeline is in fabrication of high-resolution microfeatures and with relatively optimal throughput, as 10-20 chips can be produced from a single wafer. However, fabrication of the models taller than 500 µm is challenging and only step-like, pseudo-3D features from a single side of the PDMS layer can be fabricated. To overcome these limitations, we explored several other complementary fabrication strategies.

Commercial 3D printing techniques such as fused deposition modeling (FDM) and stereolithography (SLA) were used for the production of larger models (Formlabs Form 4). However, these methods lacked the resolution and surface finish required for high-quality PDMS molding and were suitable only for some early-stage proof-of-concept prototypes.

Instead, here we employed 2PP lithography using the UpNano NanoOne printer to fabricate detailed 3D features of the chip. In brief, 3D models of the microchip were created in Autodesk Fusion, exported as STL files, and processed using UpNano Think3D software. The models were printed on glass coverslips using UpPhoto resin (UpNano). NanoOne stands out as the most developed printer for mesoscale manufacturing, enabling printing of large structures (10-20 mm in XY and up to 40 mm height) due to several advanced technologies: a submerged printing mode in a large resin reservoir, high-power 1000 W, 780 nm laser for fast printing with 5X 0.25 NA objective and adaptive resolution technology that significantly reduces print times (up to 450 mm3/h). After printing, the coverslips were washed in isopropanol and silanized overnight using vapor-phase trichloro(1H,1H,2H,2H-perfluorooctyl) silane (Sigma-Aldrich). PDMS was cast, degassed, cured at 80°C and then replicas were cut and plasma-bonded to glass-bottom dishes. This method allowed us to fabricate high-quality prototypes for validation of the 3D microstructural design elements. However, the main limitation is the upscaling technology for fabrication of the large number of chips. More importantly, process flow is still resulting in the 3D topography features only from one side of the PDMS replica.

To overcome the latter problem, commonly some labs fabricate multi-layered PDSM chips. In this approach, individual PDMS layers are fabricated separately, aligned, and plasma-bonded together. This direction was explored early on for fabrication of a few prototypes, however the alignment process is very tedious and not suitable for large-scale chips manufacturing with the acceptable quality.

To address this, inspired by industrial injection molding technology, we developed a cassette mold for PDMS curing. Molds were manufactured by CNC milling of acrylic, aluminum, or steel, depending on the desired feature size and quality. Each cassette consisted of two mold parts bearing microtopographies of the chip’s bottom and top surfaces. When assembled, the parts fit precisely, enabling accurate alignment of both sides of the device. PDMS was injected into center space of the mold, degassed in a vacuum chamber and cured in an 80°C oven. After curing, the mold was opened and the PDMS sheets were cut and bonded to glass-bottom dishes. This process overcomes many of the limitations of previously described techniques and enables scalable production of hundreds of high-quality chips in a single batch. The main drawback is the need to adapt design features for CNC milling — some smallest features must be enlarged and sharp corners rounded. However, we didn’t consider this as a drawback as concurrently we began exploring industrial plastic molding technologies to eliminate the last remaining problem: the PDMS material itself. Although PDMS is ideal for prototyping, its high absorption of small molecules makes it unsuitable for translational applications and drug testing. Many of the learnings and design optimizations from the PDMS cassette-molding process were directly translated into the design of the final injection-molded plastic chip. Currently, the chip body is fabricated from cyclic olefin copolymer (COC) due to its low shrinkage, chemical resistance, and optical transmission. The main body is fabricated as a single piece, followed by bonding of a 170 µm COC film to the bottom surface. A matching lid converting the slide was later designed and produced separately also via COC injection molding process. After assembly, chips are packaged and sterilized using ethylene oxide gas (EOG). Large-scale chip production is outsourced to an external manufacturing facility.

### Stamps fabrication

Stamps featuring crypt-mimicking microtopography were fabricated from PDMS via replica molding of 2PP-printed models. First, arrays of colon crypt-mimicking micropillars were designed in Autodesk Fusion. Several topographies were tested and optimized, the final microtopography design consists of the slightly conical pillars, ranging from 70 to 120µm in diameter (top to bottom) and 700µm in height. The 3D model was printed using a NanoOne (UpNano) 2PP printer on glass coverslips with UpPhoto resin and a 10X 0.4 NA objective.

To generate a master mold with inverted topography, the printed structures were plasma-activated and silanized overnight using vapor-phase trichloro(1H,1H,2H,2H-perfluorooctyl) silane (Sigma-Aldrich). PDMS (Sylgard 184, Dow Corning; 10:1 base-to-curing agent ratio) was poured over the master, degassed under vacuum, and cured at 80°C for 12h. The resulting PDMS replica was cut, plasma-activated, and silanized overnight as above. This PDMS negative was then used repeatedly as a mold for casting final PDMS stamps using the same procedure (Sylgard 184, 10:1 ratio, cured at 80°C for 12h). Microstamping hydrogel topography

To generate the extracellular matrix (ECM) for the chip bovine TeloCol-6 Type I Collagen solution (5225-50mL, Advanced Biomatrix) was neutralized following manufacturer instructions and then mixed with Matrigel (356231, Corning) in 75/25 (v/v) ratio. A PDMS stamp with crypt-mimmicking array of pillar topography was placed in the central top reservoir. After gel injection through gel-loading port, hydrogel was polymerized at 37°C, 5% CO_2_ (hereafter referred to as “incubator”) for 15 min. After ECM polymerization, all chip reservoirs were topped up with PBS or cell culture media. Prepared chips with hydrogels can be stored for multiple weeks in the incubator or in the +4°C fridge before use.

### Seeding and culture of engineered intestine

For seeding of the chips - hereafter referred to as ‘engineered intestines’ once intestinal epithelial cells are included - approximately 4-day-old human intestinal organoids from various regions of the gastrointestinal tract (duodenum, jejunum, ileum, colon) between passage 5-25 were collected in base-media (Supplementary Table 1) and transferred to a 15 mL Falcon tube. Following centrifugation at 300g at 4°C for 4 min, the base-media (Supplementary Table 1) was aspirated and the pellet was thoroughly resuspended in base-media. After repeating the centrifugation step, organoids were singularized with TrypLE express solution (12604-013, Gibco) containing 250 U/mL DNase I (4536282001, Roche) and 10 µm Y-27632 (72305, Stemcell Technologies) for 8 min at 37°C before harsh resuspension. This step was performed twice before swiftly blocking the dissociation by adding base-media with 10% FBS (Supplementary Table 1). Next, cells were passed through a 40 µm large number of chips. More importantly, process flow is still resulting in the 3D topography features only from one side of the PDMS replica.

To overcome the latter problem, commonly some labs fabricate multi-layered PDSM chips. In this approach, individual PDMS layers are fabricated separately, aligned, and plasma-bonded together. This direction was explored early on for fabrication of a few prototypes, however the alignment process is very tedious and not suitable for large-scale chips manufacturing with the acceptable quality.

To address this, inspired by industrial injection molding technology, we developed a cassette mold for PDMS curing. Molds were manufactured by CNC milling of acrylic, aluminum, or steel, depending on the desired feature size and quality. Each cassette consisted of two mold parts bearing microtopographies of the chip’s bottom and top surfaces. When assembled, the parts fit precisely, enabling accurate alignment of both sides of the device. PDMS was injected into center space of the mold, degassed in a vacuum chamber and cured in an 80°C oven. After curing, the mold was opened and the PDMS sheets were cut and bonded to glass-bottom dishes. This process overcomes many of the limitations of previously described techniques and enables scalable production of hundreds of high-quality chips in a single batch. The main drawback is the need to adapt design features for CNC milling — some smallest features must be enlarged and sharp corners rounded. However, we didn’t consider this as a drawback as concurrently we began exploring industrial plastic molding technologies to eliminate the last remaining problem: the PDMS material itself. Although PDMS is ideal for prototyping, its high absorption of small molecules makes it unsuitable for translational applications and drug testing. Many of the learnings and design optimizations from the PDMS cassette-molding process were directly translated into the design of the final injection-molded plastic chip. Currently, the chip body is fabricated from cyclic olefin copolymer (COC) due to its low shrinkage, chemical resistance, and optical transmission. The main body is fabricated as a single piece, followed by bonding of a 170 µm COC film to the bottom surface. A matching lid converting the slide was later designed and produced separately also via COC injection molding process. After assembly, chips are packaged and sterilized using ethylene oxide gas (EOG). Large-scale chip production is outsourced to an external manufacturing facility.

### Stamps fabrication

Stamps featuring crypt-mimicking microtopography were fabricated from PDMS via replica molding of 2PP-printed models. First, arrays of colon crypt-mimicking micropillars were designed in Autodesk Fusion. Several topographies were tested and optimized, the final microtopography design consists of the slightly conical pillars, ranging from 70 to 120µm in diameter (top to bottom) and 700µm in height. The 3D model was printed using a NanoOne (UpNano) 2PP printer on glass coverslips with UpPhoto resin and a 10X 0.4 NA objective.

To generate a master mold with inverted topography, the printed structures were plasma-activated and silanized overnight using vapor-phase trichloro(1H,1H,2H,2H-perfluorooctyl) silane (Sigma-Aldrich). PDMS (Sylgard 184, Dow Corning; 10:1 base-to-curing agent ratio) was poured over the master, degassed under vacuum, and cured at 80°C for 12h. The resulting PDMS replica was cut, plasma-activated, and silanized overnight as above. This PDMS negative was then used repeatedly as a mold for casting final PDMS stamps using the same procedure (Sylgard 184, 10:1 ratio, cured at 80°C for 12h). Microstamping hydrogel topography

To generate the extracellular matrix (ECM) for the chip bovine TeloCol-6 Type I Collagen solution (5225-50mL, Advanced Biomatrix) was neutralized following manufacturer instructions and then mixed with Matrigel (356231, Corning) in 75/25 (v/v) ratio. A PDMS stamp with crypt-mimmicking array of pillar topography was placed in the central top reservoir. After gel injection through gel-loading port, hydrogel was polymerized at 37°C, 5% CO_2_ (hereafter referred to as “incubator”) for 15 min. After ECM polymerization, all chip reservoirs were topped up with PBS or cell culture media. Prepared chips with hydrogels can be stored for multiple weeks in the incubator or in the +4°C fridge before use.

### Seeding and culture of engineered intestine

For seeding of the chips - hereafter referred to as ‘engineered intestines’ once intestinal epithelial cells are included - approximately 4-day-old human intestinal organoids from various regions of the gastrointestinal tract (duodenum, jejunum, ileum, colon) between passage 5-25 were collected in base-media (Supplementary Table 1) and transferred to a 15 mL Falcon tube. Following centrifugation at 300g at 4°C for 4 min, the base-media (Supplementary Table 1) was aspirated and the pellet was thoroughly resuspended in base-media. After repeating the centrifugation step, organoids were singularized with TrypLE express solution (12604-013, Gibco) containing 250 U/mL DNase I (4536282001, Roche) and 10 µm Y-27632 (72305, Stemcell Technologies) for 8 min at 37°C before harsh resuspension. This step was performed twice before swiftly blocking the dissociation by adding base-media with 10% FBS (Supplementary Table 1). Next, cells were passed through a 40 µm strainer (43-50040-51, PluriSelect) before centrifugation at 500 g at 4°C for 4 min. Afterwards, the cells were resuspended in chip-media without A-83 (Supplementary Table 1) containing 10 µm Y-27632. Approximately 1.5x105 cells were seeded in a volume of 2.5 µl onto each micropatterned hydrogel. 150 µl chip-media (- A-83, + 10 µm Y-27632) was immediately added to each side reservoir of the engineered intestines, while the mid chamber remained empty to allow the cells to attach to the ECM. After 30 min in the incubator, 100 µl chip-media (- A-83, + 10 µm Y-27632) was carefully added to the mid chamber to avoid cell detachment. Culture medium was exchanged every second day, with the removal of EGF and addition of A-83 on day 4, after full monolayer formation (Supplementary Table 1).

### Immune cell hydrogel seeding and culture of immunocompetent engineered intestines

On the day of hydrogel seeding, TRMs or PBMCs were thawed following our previously described protocol^133^. T cells were isolated from PBMCs using the EasySep™ Human T Cell Isolation kit (17951, STEMCELL Technologies) and EasySep™ Magnet (18000, STEMCELL Technologies) following manufacturer instructions. Viability and number of immune cells was assessed with Trypan Blue (216040, Invitrogen) on a Countess II (Invitrogen). If the viability of the thawed TRM crawl-out was < 75% (due to remaining tissue debris), Dead Cell Removal Kit (130-090-101, Miltenyi Biotec) was applied with MS columns (130-042-201, Miltenyi Biotec) on the OctoMACS™ separator (130-042-109, Miltenyi Biotec) following manufacturer instructions (typically improved to > 85% viability). Afterwards, immune cells were placed in the incubator until further processing. Chips were prepared as described above until hydrogel loading. Then, ∼1.3x105 immune cells were mixed with 6 µL of ECM per chip. Next, 22 µL of ECM was dispensed through the loading port per chip into the mid reservoir. Immediately after, 6 µL of the ECM-immune suspension was carefully dispensed onto a flipped PDMS stamp, with the micropattern facing upward, and transferred using forceps onto the previously-dispensed ECM in the mid reservoir of the engineered intestine. Next, immune-hydrogel suspensions were polymerized at 37°C for 15 min. After polymerization of the ECM, the immune-chip-slide was topped up with immune-chip-media (Supplementary Table 1) to ease microstamp removal. After organoid dissociation and epithelial cell seeding as described above, 150 µl immune-chip-media (- A-83, + 500 pg/mL IL-15 (570306, BioLegend), + 10 µm Y-27632) was immediately added to each side reservoir of the engineered intestine, while the mid chamber remained empty to allow for cells to attach to the ECM of the engineered intestine. After 30 min in the incubator, 100 µl immune-chip-media (- A-83, + 500 pg/mL IL-15, + 10 µm Y-27632) was carefully added to the mid chamber. Immune-chip-media (+ 500 pg/mL IL-15) was exchanged every second day, with the removal of EGF and addition of A-83 on day 4, after full monolayer formation.

### Permeability assessment of engineered intestinal epithelium

To assess the integrity of the engineered intestinal epithelium in homeostatic conditions, the apical and basal media were aspirated thoroughly. Next, 120 µl of a 100 µg/mL solution of Cascade Blue hydrazide, trilithium salt (C3239, Invitrogen) in chip-media (Supplementary Table 1) were added to the apical reservoir. 170 µl of chip-media without Cascade Blue were added to the side reservoirs. The assay was carried out for 180 min in the incubator. Subsequently, the fluorescence of the basal medium was quantified using Cell Culture Microplates (655090, Greiner Bio-one) in the Flex Station 3 plate reader (Molecular Devices), with measurements taken from top at an excitation wavelength of 380 nm and an emission wavelength of 420 nm. The concentration of Cascade Blue that diffused to the basal compartment was calculated against a standard curve and the resulting apparent permeability (P_app_) was quantified using the following formula:

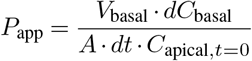

where *V* _basal_ is the volume of the basal compartment, *dC*_basal_ is the measured concentration in the basal compartment, *A* is the area of the epithelial monolayer grown on the ECM (0.19477 cm^2^), *dt* is the time of the assay and *C*_apical, t=0_ is the initial concentration added to the apical reservoir at the start of the assay. For infection experiments, one gentle PBS wash was performed in the apical reservoir of control and infected samples to remove the majority of the grown biofilm prior permeability assessment. The Cascade Blue, in addition to 200 µg/mL of gentamicin (G1397-10ML, Sigma-Aldrich), were supplemented in the corresponding medium of the experiment. The assay was conducted over a period of 90 min, and fluorescence was measured from the bottom using a Cell Culture Microplate that was covered with a transparent adhesive foil on top.

### Mucus accumulation strategy and staining

Secreted mucus was accumulated by reducing the total apical volume from ∼120-100 µl of chip-media to either ∼30 µl or ∼5 µl, based on the experimental needs. Generally, ∼30 µl allowed for an accumulation of a less dense mucus that stayed hydrated. Accumulation with ∼5 µl allowed for a denser layer of mucus that temporarily dried during the day and acquired a yellow hue. The mucus accumulation strategy was initiated on day 4-5 of culture after full coverage of the ECM surface and appearance of goblet cells. The accumulation strategy was continued for 4-5 days and the accumulation of mucus on the epithelium surface was monitored by bright field microscopy. In the accumulation with ∼5 µl, fresh 5 µl of chip-media was added every day due to the liquid absorption by the intestinal epithelium and resulting air-liquid interface at the center of the apical reservoir. In the accumulation with ∼30 µl, the volume was monitored every day and adjusted to ∼30 µl in case of detected liquid release from the epithelium or liquid absorption or air-liquid interface. The whole volume was exchanged every second day.

Staining of the glycocalyx and secreted mucus in adult and developing engineered epithelia was performed by an overnight incubation with 30 µl or 5 µl of 488 Wheat Germ Agglutinin (WGA) (25 µg/mL, W11261, Invitrogen), respective of the accumulation strategy. In cases where higher fluorescence signal of the secreted mucus was desired, additional WGA incubations were performed in consecutive days. Shed cells were stained with an overnight incubation with SiR700 (3µM, SC015, Sphirochrome) in chip-media. Confocal images were acquired as described in Methods: Confocal microscopy imaging. 3D reconstruction of the mucus was generated using the Leica Application Suite X (LAS X) software (4.6.1.27508) and adjusted for visualization purposes.

### Proteomic assessment of the luminal content

For proteomic assessment of the luminal content, the mucus of the engineered intestines was accumulated for 5 days, reducing the luminal volume from ∼100 µl to ∼5 µl. In these samples, DMEM/ F12 (31331-028, Gibco) + HEPES (10 mM) + Primocin (50 µg/mL) was used as apical medium. For collection, the luminal content was collected by pipetting 5 µl of fresh apical medium up and down thoroughly and transferred to a Protein LoBind 1.5 mL tube (022431081, Eppendorf). This step was repeated a total of four times. The samples were immediately stored at - 80°C until further use.

Samples were processed into peptides using the iST kit (PreOmics), according to the manufacturer’s instructions. Mass spectrometry analysis was conducted on a Orbitrap Fusion™ Lumos™ Tribrid™ Mass Spectrometer (ThermoFisher Scientific) device equipped with a Nanospray Flex Series Ion Source and a PRSO-V2 sonation oven (Sonation), which was coupled to an EASY-nLC™ 1200 system (ThermoFisher Scientific). Samples were resuspended in 2% acetonitrile, 0.5% formic acid in water and loaded onto a C18 trap column (Acclaim™ PepMap™ 100 C18 HPLC Columns 20 mm x 75 µm; 3µm) at a flow rate of 20 µl/ min, peptides were separated using a 75-µm inner diameter C18 column (Aurora Column 25 cm x 75 µm C18 1.7 µm; Gen3; (IonOpticks). Chromatographic separation was achieved using the following conditions: Buffer A: 0.1% formic acid in water; Buffer B: 80% 0.1% formic acid in acetonitrile. The column oven and autosampler tray were held at 40°C and 7°C, respectively. The chromatographic gradient was run at a flow rate of 250 nL/min as follows: 0-47 min: linear gradient from 9 to 20% B; 47-90 min: linear gradient from 20 to 40% B; 90-92 min: linear gradient from 40 to 97 %; 92 to 104 min: hold at 97%. The mass spectrometer was operated in positive mode with the spray voltage set to 1.7kV and the ion transfer tube held at 275 °C. All measurements were performed in data-dependent mode. Precursor mass spectra were recorded in a range of 375-1500 m/z with the resolution set at 240,000 the AGC target set in the standard mode and the maximum injection time in auto mode. For MS/MS acquisition, data recorded in centroid mode with injection time was set to maximum 50 milliseconds with a Gain Control setup at 100%. Normalized collision energy was set to 30. All experiments were recorded in ITMS-mode, scan rate was turbo. Data was extracted and searched against the SwissProt database (Jun 2024) by using Proteome Discoverer 2.5 (Thermo Scientific) and Mascot version 2.8.2 (Matrix Science). All cysteines were considered alkylated (carbamidomethyl) and oxidation of methionine, and protein N-terminal acetylation were allowed as variable modifications. Data were first searched using fully tryptic constraints. Matched peptides were filtered using a Percolator-based 1% false discovery rate.

iBAQ (Intensity-based absolute quantification) values were calculated with an in-house R script using R (4.3.0), Biostrings (2.68.1), and tidyverse (2.0.0). In brief: the number of theoretically possible tryptic peptides with 6 to 30 amino acids was calculated based on an in-silico tryptic digest of the human SwissProt database already used for the Proteome Discoverer (PD) search. The theoretical peptide count was then mapped to the PD output by protein accession. Finally, the iBAQ value was calculated by dividing the respective protein abundance by the theoretical number of tryptic peptides. Proteins with only one unique peptide and any missing values for any of the engineered intestine replicates within each donor were excluded from the analysis. IBAQ values for each protein were averaged first across engineered intestines within each donor and then across donors to generate the final IBAQ rank plots.

### Commensal bacteria co-culture experiments

Enterococcus faecium (19434, ATCC) inoculums were grown overnight at 230 rpm, 37°C in 4 mL of Brain Heart Infusion (BHI; 1002417, MP Biomedicals) medium. Lacticaseibacillus rhamnosus (7469, ATCC) inoculums were grown overnight at 230 rpm, 37°C, 5% CO_2_, and 95% humidity in 10 mL of De Man, Rogosa and Sharpe (MRS) medium (1106610500, Millipore). For both strains, 50 mL Mini Bioreactor tubes (431720, Corning) were used for the overnight cultures. Following overnight growth, the OD_600_ values of the cultures were measured to adjust the bacterial concentration to 10^8^ CFU/mL. The bacteria were then pelleted by centrifugation at 10000 rpm (*E. faecium*) or 5000 rpm (*L. rhamnosus*) for 1 min, and washed once with PBS. Subsequently, the pellets were resuspended in 1 mL of DMEM, low glucose, pyruvate (31885-023, Gibco) + HEPES (1:100, 15630080, Gibco) medium and serially diluted. In the case of *L. rhamnosus*, a 10% (v/v) supplementation of MRS was added. Finally, the desired number of bacteria was added in 30 µl of the corresponding medium to the apical compartment of the engineered intestines, which were cultured in the absence of primocin since day 0. The co-cultures were maintained in the incubator until the end of the experiment. To measure the number of bacteria on the samples, the luminal content was pipetted up and down thoroughly, and transferred to a 1.5 mL tube. This step was repeated with extra 100 µl PBS washes. A serial dilution of the total collected content was performed and the bacteria were plated and counted in TSA plates (43019, Biomerieux).

### *S*.Tm culture and infection experiments

The GFP reporter strain of *Salmonella enterica* subsp. enterica serovar Typhimurium (14028GFP, ATCC), hereafter referred to as *S*.Tm, was used to infect the epithelial monolayer in the presence and absence of TRMs. In all cases, *S*.Tm inoculums were grown overnight at 37°C and 230 rpm in 10 mL of BHI (1002417, MP Biomedicals) medium with 100 µg/mL ampicillin (A53542, Sigma-Aldrich) using 50 mL Mini Bioreactor tubes (431720, Corning). Following overnight growth, the OD_600_ values of the cultures were measured to adjust the bacterial concentration to 10^8^ CFU/mL. The bacteria were then pelleted by centrifugation at 3000 rpm for 5 min and washed once with PBS. Subsequently, the pellets were resuspended in 1 mL of the corresponding apical medium (see next) and serially diluted. The desired number of bacteria was then added to the apical compartment of the engineered intestines for infection experiments.

#### Epithelium-*S*.Tm co-culture experiments

For the epithelium-*S*.Tm infection experiments, the engineered intestines were cultured in chip-media (Supplementary Table 1) without primocin until the day of infection. Different mucus accumulation strategies were followed depending on the condition (Mucus^Hi^: 4 days, 5 µl, Mucus^Int^: 2 days, 5 µl, Mucus^Lo^: 4 days, 120 µl and 1x wash prior infection). On the day of infection, 10^5^ CFU of *S*.Tm were resuspended in chip-media (- primocin) and added to the apical compartment in a total volume of 5 µl to preserve the dense mucus structure. The infection was maintained in the incubator until the end of the experiment (18 h or 24 h). To measure the number of bacteria on the samples, the luminal content was resuspended in 100 µl of PBS in-situ, pipetted up and down thoroughly, and transferred to a 1.5 mL tube. This step was repeated one more time. A serial dilution of the collected content was performed and the bacteria were plated and counted in TSA plates (43019, Biomerieux).

#### TRM-Epithelium-*S*.Tm tri-culture experiments

For the TRM-epithelium-*S*.Tm infection experiments, the engineered intestines were cultured in immune-chip-media without primocin until the day of infection. On the day of infection, following the addition of basal immune-bacteria-chip-media (+ 10 µl/mL gentamicin; Supplementary Table 1) to the basal reservoirs of the engineered intestines, 10^2^ CFU of *S*.Tm were resuspended in apical immune-bacteria-chip-media (Supplementary Table 1) and added to the apical compartment in a total volume of 100 µl. Mucus was not accumulated in these samples. The infection was maintained in the incubator until the end of the experiment. To measure the number of bacteria on the sample, the apical volume was collected with gentle pipetting and transferred to a 1.5 mL tube. This step was repeated with 100 µl of PBS and the total volume content was serially diluted, plated and counted in TSA plates.

### Gentamicin protection assay

Gentamicin protection assays were carried out using duodenal epithelia at day 10 of culture. *S*.Tm inoculums were grown for 12 h in LB (10855021, Gibco) + 0.3M of NaCl (S/3161/60, Thermo Fisher Scientific) + 100 µg/mL ampicillin (A53542, Sigma-Aldrich) at 230 rpm, 37°C. Next, the cultures were diluted 1:20 and incubated for 3 h in the same conditions. ∼10^7^ CFU were added to each sample and co-cultured for 2 h 30 min in the incubator. After that time, the apical compartment was washed twice with DPBS, and 200 µg/mL of gentamicin in chip-media was added to the apical and basal compartments. The engineered intestines were incubated for 18 h, after which the epithelium was collected with tweezers and transferred to a tube containing 100 µl of 1% Triton X-100 (Sigma-Aldrich, T8787). The content was incubated for 15 min and pipetted up and down at 0 min, 7 min and 15 min. A serial dilution of the lysate was plated and counted in TSA plates.

### Legendplex cytokine analysis of engineered intestines

To analyze the cytokines in the supernatant of the engineered intestines, 100 µL supernatant was collected from both basal reservoirs per engineered intestine and pooled afterwards. Supernatants were quickly centrifuged to remove cell debris and frozen at -20°C, followed by a transfer to -80°C until further processing. Assessment of the cytokines was performed using the LEGENDplex Human CD8/NK Panel (741187, BioLegend) and a custom designed LEGENDplex panel (CXCL10, IL-8, Granzyme B, IFNγ, IL-1β, IL-2, IL-6, IL-10, CCL2, sFasL, sFas, TGF-β1 (free active), TNF-; BioLegend). Controls, standards and beads were prepared following the manufacturer instructions. Samples were not diluted. Briefly, 25 uL of the samples were mixed with 25 µL Assay buffer and 25 µL mixed beads. The 25 µL diluted standard was mixed with 25 µL assay buffer and 25 µL mixed beads. Both samples and standard mixtures were dispensed in a V-bottom plate. Afterwards, samples were incubated for 2 hours at RT on a shaker at 800 rpm. Samples were centrifuged at 500g for 2 min followed by removal of the supernatant. Samples were washed with 200 µL 1X wash buffer per well, followed by addition of 25 µL detection antibodies and incubation for 1h at RT on a shaker at 800 rpm. Next, without washing, SA-PE was added and incubated for 30 min at RT on a shaker at 800 rpm. Samples were centrifuged at 500g for 2 min followed by removal of the supernatant to remove excess antibodies. Lastly, 150 µL of 1X wash buffer was added before acquisition on Cytek Aurora with the 5L configuration (Cytek) following the recommended experimental setup. To obtain absolute cytokine concentrations, the raw data were processed and analyzed using the software provided by the manufacturer (LEGENDplex data analysis software, BioLegend).

### Confocal microscopy imaging

The following microscopes were used for confocal imaging of the stained and live engineered intestines:

1. Nikon Ti2 microscope with CSU-W1 spinning disk (Yokogawa) equipped with 20XC S Plan Fluor LWD, 25X Apo LWD 1.1W DIC N2, sCOMS camera (ORCA-Fusion BT, Hamamatsu Photonics). Depending on the application the following objectives were used: 4X PLAN APO Lambda OFN25, 10X PLAN APO Lambda OFN25 DIC N1, 25XC Sil Plan Apo Lambda S (Nikon Instruments). The Extended Depth of Focus (EDF) feature of the NIS-Elements software (Nikon) was used to create focused bright field and fluorescent images of the engineered intestines. In certain cases, images were further processed and color adjusted using ImageJ software (2.3.0/1.54f).
2. Leica STELLARIS 8 confocal microscope equipped with 3 Power HyD detectors, White Light Laser (440 nm - 790 nm) using 25X HC FLUOTAR L VISIR X25 objective (Leica Microsystems). Images were either visualized as maximum projection or three-dimensionally reconstructed using the Leica Application Suite X (LAS X) software (4.6.1.27508). Imaris 3D reconstructions: 3D reconstructions of the epithelial layer with integrated TRMs from the engineered intestines whole mount stainings were generated with the Imaris surface function (Version 9.9.1). Surfaces were rendered based on fluorescence intensities of the iCas9 CDH1-mScarlet3 reporter and CD3, CD8 antibody stainings.

### Time-lapse imaging of engineered intestines

Prior to immune cell seeding, TRMs or T cells derived from PBMCs (isolation described above) were labelled with Cell Trace Far Red (C34572, Invitrogen) following manufacturer instructions. Afterwards, immuno-chips were prepared as described above. Time-lapse imaging was performed 7-10 days (where indicated) post co-culture setup with either a STELLARIS 8 confocal microscope with a 25X HC FLUOTAR L VISIR X25 objective (Leica Microsystems) or with Nikon Ti2 microscope equipped with CSU-W1 spinning disk (Yokogawa) using with 25X Apo LWD 1.1W DIC N2 objective (Nikon Instruments). Images on Leica STELLARIS 8 were obtained in bidirectional mode with 1024 x 1024 pixels at 600 Hz. Images were acquired with varying z-stack depth (z-steps, 4 µm). Samples were imaged between 42 and 60 min (where indicated). During imaging, samples were maintained in a microscopy incubation chamber 37°C, 5% CO_2_ (The Box, Life Imaging Services). For live-imaging along the crypt-villus axis, engineered intestines were collected from the glass-slide and cut in two followed by placing the engineered intestines perpendicularly on top of a glass-dish. The engineered intestines were then fixated with a thin glass coverslip to avoid buoyancy due to media in the dish. After acquisition, maximum-intensity projections were generated for downstream analysis.

Time-lapse movies were analysed using CellProfiler. Briefly, cells were segmented at each frame with multiotsu, morphological parameters were extracted and segmented cells were tracked over time using LAP algorithm with following parameters: Number of Standard Deviations for search radius=80, search radius limits= 3px-80px, Gap closing cost=2, Split Alternative cost=1, Merge Alternative cost=1, Mitosis alternative=1, Maximum gap displacement=100px, Maximum split score=10, maximum merge score=5, maximal temporal gap=6, maximum mitosis distance=40. The output per cell was analysed using KNIME. The means and standard deviation of morphological features per track over time were calculated. The standard deviation of morphological features over time is a measure of the dynamism of cells as they use amoeboid motility. Additionally, track life time and track total distance were calculated. Tracks with less than 5 frames were discarded.

### Whole-mount staining

For fixed whole-mount staining and imaging, engineered intestines were washed once with 1X DPBS before fixation with 4% paraformaldehyde (PFA) in the chip-slide. After 1 hour of fixation at RT, the engineered intestines were washed three times with 1X DPBS. Engineered intestines were either stained in-situ or collected using a biopsy puncher (3 mm diameter) and transferred into a 24-well plate. Next, samples were permeabilized with 0.2% Triton X-100 (Sigma-Aldrich, T8787) in 1X DPBS for 2 h at RT on an orbital rocker at 50 rpm. Permeabilization was followed by a blocking step with blocking buffer (10% donkey serum (S30-M, Sigma-Aldrich), 0.01% Triton X-100 in DPBS 1X) for 4 h at RT on an orbital rocker or 16 h at 4°C. Primary antibodies (Supplementary Table 1) were applied and incubated overnight at 4°C on an orbital rocker.

Afterwards, samples were washed three times for 1 h each at RT with 1X DBPS on an orbital rocker. Anti-species secondary antibodies (Supplementary Table 1), DAPI (1 µg/mL, 62249, Thermo Scientific), and phalloidin (Alexa Fluor 568 Phalloidin A12380; Alexa Fluor 488 Phalloidin A12379; Alexa Fluor 647 Phalloidin A30107, Thermo Fischer Scientific) were incubated for 4 h at RT or 16 h at 4°C on an orbital rocker, covering the sample with aluminum foil. Samples were washed three times for 1 h each with DPBS on an orbital rocker prior imaging.

### FFPE embedding of engineered intestines

#### Regular workflow

Engineered intestines were washed once with 1X DPBS before fixation with 4% paraformaldehyde (PFA) in the chip-slide. After 1 hour of fixation at RT, the engineered intestines were washed three times before using a biopsy puncher (3 mm diameter) to detach the ECM and tissue. In parallel, the engineered intestines histomold, specifically designed to facilitate co-planar embedding of up to 50 engineered intestines, was prepared as previously described^17^. Briefly, liquid histogel was dispensed into the chip-histomold and allowed to polymerize for 20 min at 4°C. Following polymerization, the histoarray was demoulded. Engineered intestines were then transferred with a thin tweezer into the slits of the histoarray. After careful glueing of the engineered intestines in the wells of the histoarray (see Harter et al.17), the histoarray was completely filled and distributed into biopsy cassettes. Samples were dehydrated overnight using a Vacuum filter processor (HistoCore PEARL, Leica). The following day, samples were embedded in liquid paraffin.

#### Workflow to retain luminal content

The basal compartments of the engineered intestines were washed once with 1X DPBS before fixation with 4% paraformaldehyde (PFA). The apical reservoir was neither washed nor fixed to preserve the integrity of the luminal content. After 2 h of fixation at RT, the engineered intestines were washed three more times only in the basal compartment. Next, the bottom glass of the chip-slide was detached by applying mechanical pressure and the matrix was punched from the bottom of the hydrogel in upward direction, keeping the epithelium and luminal content. The punched sample was then carefully covered with 8 µL of TeloCol®-6 Type I Collagen solution (5225-50mL, Advanced Biomatrix) neutralized according to manufacturer instructions. Finally, the sample was incubated for 15 min at 37°C before the transfer to the slits of the histoarray, as described before.

### Microtome sectioning

FFPE blocks were, unless otherwise specified, sectioned at a thickness of 3.5 µm and transferred on Superfrost Ultra Plus Gold Adhesion Slides. Slides were incubated in a slide oven overnight at 37°C.

#### H&E and AlcianBlue/ Periodic Acid–Schiff staining and imaging

H&E staining was executed in a fully automated manner following the standard protocol on a Ventana HE600 stainer (Roche Tissue Diagnostics). AlcianBlue/ PAS staining was performed fully automated using the AlcianBlue for PAS (860-003, Ventana) and PAS staining (860-014, Ventana) kits on the Ventana Benchmark (Roche Tissue Diagnostics). Briefly, after three cycles of deparaffinization, slides were washed and 200 µL of PAS AlcianBlue incubated for 16 min at 37°C. Wash was repeated, and 200 µL of PAS periodic acid was applied for 4 min, before another wash and 12 min of PAS Schiffs staining. Next, slides were washed again and 200 µL PAS neutralizer was added and incubated for 4 min. Lastly, slides were washed with 100 µL before counterstaining the slide with 200 µL of PAS hematoxylin for 8 min. Afterwards, slides were coverslipped as indicated in the manufacturer instructions. H&E, IHC and AlcianBlue/ PAS stained slides were imaged with a brightfield whole-slide scanner at X40 (Hamamatsu, NanoZoomer, S360; pixel size 0.23 µm/px).

### Immunohistochemistry staining

As previously described^133^, IHC staining of FFPE slides was performed using a Ventana Discovery Ultra automated tissue stainer (Roche Tissue Diagnostics). The slides were baked for 8 minutes at 60°C, followed by a deparaffinization cycle for 8 min at 69°C (deparaffinization cycle repeated three times). Heat-induced antigen retrieval was performed with Tris-EDTA buffer (pH 7.8, CC1, 950-227, Ventana) at 95°C for a total of 40 min. Depending on the host species of secondary antibody, the slide was incubated with either DISCOVERY Goat Ig Block (760-6008, Ventana) or 10% donkey serum for 32 min at 37°C followed by application of DISCOVERY Inhibitor (760-4840, Ventana) for 8 min. Afterwards, primary antibodies were applied (diluted in Discovery Ab diluent, 760-108, Ventana; for specific concentrations and incubation times see Supplementary Table 1). Primary antibodies were detected using anti-species secondary antibodies conjugated with horseradish peroxidase (HRP; OmniMap anti-rabbit HRP, 760-4311, Ventana; OmniMap anti-mouse HRP, 760-4310, Ventana; OmniMap anti-rat HRP, 760-4310, Ventana; OmniMap anti-goat HRP, 760-4647, Ventana) for 16 min and visualized by conversion of 3,3’-diaminobenizidine (DAB; Discovery ChromoMap DAB kit, 760-159, Ventana). Samples were counterstained with haematoxylin (Haematoxylin II, 790-2208, Ventana) and blueing reagent (760-2037, Ventana). After dehydration with a standard series of alcohol (75%, 95%, 100%, 100% v/v; CAS64-17-5, Roche) followed by 100% xylol baths (444240050, ACROS Organoids), slides were coverslipped in the RCM7000 coverslipper (MEDITE) using a standard histology glue (00811-EX, Pertex). Slides were dried for at least 2 h before imaging.

### FFPE-based mIF

As previously described^133^, Opal mIF staining of FFPE slides was performed using a Ventana Discovery Ultra automated tissue stainer (Roche Tissue Diagnostics). The slides were baked for 8 minutes at 60°C, followed by a deparaffinization cycle for 8 min at 69°C (deparaffinization cycle repeated three times). Heat-induced antigen retrieval was performed with Tris-EDTA buffer (pH 7.8, CC1, 950-227, Ventana) at 95°C for a total of 40 min. Depending on the host species of secondary antibody, the slide was incubated with either DISCOVERY Goat Ig Block (760-6008, Ventana) or 10% donkey serum for 32 min at 37°C followed by application of DISCOVERY Inhibitor (760-4840, Ventana) for 8 min. Afterwards, primary antibodies were dispensed onto the slide (diluted in Discovery Ab diluent, 760-108, Ventana; for specific concentrations and incubation times see Supplementary Table 1). Primary antibodies were detected using anti-species secondary antibodies conjugated with HRP (see above for catalogue numbers) for 16 min. Next, the corresponding Opal dye (Opal 480 FP150001KT, Opal 520 FP1487001KT, Opal 570 FP1488001KT, Opal 620 FP1495001KT, Opal 690 FP1497001KT, Opal 780 FP1501001KT, Akoya Biosciences) were applied, previously prepared following manufacturers instructions. After each application of primary, followed by the corresponding secondary antibody and opal dye (with increasing fluorophore excitation wavelength), an antibody neutralization and denaturation step was applied to remove residual antibodies and HRP, before starting the staining cycle again with the Discovery Inhibitor blocking step. Lastly, samples were counterstained with 4’,6-diamidino-2-phenylindol (DAPI, Roche). When used, fluorescein-labeled Jacalin (FL-1151-5, Vector Laboratories) was integrated into the mIF staining protocol by swapping it with the 520 opal dye and always applied last in the staining protocol.

### FFPE-based mIF imaging

As previously described^133^, Opal mIF stainings were imaged and digitized using multispectral imaging of the Vectra Polaris (Perkin Elmer) employing the MOTiF technology at X20 magnification for all seven Opal dyes (Opal 480-780) and DAPI. Slides were imaged in a batch manner to ensure identical imaging settings and cross-comparability within each experiment for subsequent image analysis. Unmixing of channels and processing of images were performed with PhenoChart (v.1.0.12) and inForm (v2.4). Images were processed as .qptiff tiles and fused in HALO (Indica Labs, v3.6.4134.396). Pixel resolution 0.5 µm / pixel.

High-resolution Opal mIF images were obtained using a STELLARIS 8 microscope (Leica) with a water-immersion objective (HC FLUOTAR L VISIR X25/0.95 numerical aperture WATER). A white-light laser (440-790 nm) was used to image all Opal dyes. Channel signals were acquired sequentially to reduce cross-bleeding. Images were obtained in bidirectional mode with 1.024 x 1.024 pixels (pixel size, 384.56 x 384.56 nm2) or 2.048 x 2.048 pixels (pixel size, 273.8 x 273.38 nm2). Images were either visualized as maximum projection or three-dimensionally reconstructed using the Leica Application Suite X (LAS X) software (4.6.1.27508).

### Image analysis of FFPE-based mIF

Image analysis of Opal mIF images was performed with HALO (Indica Labs, v3.6.4134.396) and HALO AI (Indica Labs, v3.6.4134). Sample annotation was either manually performed or by deploying the TMA module in HALO. For detection and annotation of the engineered intestines, a RandomForest (v2) classifier was used to distinguish organoids from background (e.g. matrix; minimum object size > 1000 µm). Following validation, engineered intestines were handled as individual regions of interest (ROI) and quantification was executed per ROI. The RandomForest (v2) classifier was integrated in the Area Quantification FL (v.2.3.4) modules and used to quantify positive marker staining against either overall size of each individual object or normalized to the DAPI+ area of the object (as indicated in legend texts) as well as for quantification of intensities. The HighPlex FL (v4.2.14) module was used to perform nuclear segmentation based on DAPI+ cells, assisted by HALO AI’s integrated ‘AI default nuclear segmentation type’) for phenotype analysis. The FISH-IF (v2.2.5) module was used to perform DAPI-based nuclear segmentation and identification of cells positive for single intracellular bacteria. Specific cell phenotypes were determined by co-localization of the respective markers of interest with the DAPI+ nuclei (including the cytoplasm radius (1.25 µm) and nuclear signals). By integrating the RandomForest (v2) or DenseNet (v2) classifier in the HighPlex analysis module, e.g. localization of the T cells in the matrix or epithelium was determined.

Assessment of the spatial patterning was performed by manually annotating the crypts and the top-epithelial layer along the E-Cadherin+ signal as separate ROIs. The basal side of the top-epithelial layer served as the ROI border to the crypt underneath. Cell quantification was then performed within the separate annotations per engineered intestine using the Highplex FL module. Similarly, quantification of the mucus thickness was performed by manually outlining two lines within the same annotation layer: one along the apical side of the E-Cadherin+ epithelium of the top-epithelial layer (excluding the crypt depth) and a second line along the lumen-facing top layer of the Jacalin+ mucus.

Positive controls on primary matched or non-matched primary intestinal tissue were included in every staining. In case of some experiments, negative controls were co-embedded within the same histoarray by experimental design, easing unbiased threshold adjustment and detection of potential unspecific antibody binding (e.g. epithelium-only control served as negative control for immune markers; different intestinal regions within one FFPE-block).

### Brightfield image analysis

Quantification of the number of viable crypts per engineered intestine was performed with a RandomForest classifier (v2) in HALO (Indica Labs, v3.6.4134.396). Dozens of viable intestinal crypts from different donors were manually annotated using brightfield images of the engineered intestines acquired with the X2 objective of the Leica Dmi1 inverted microscope (11526227, Leica Microsystems). After brief verification, the algorithm was deployed to detect the number of viable crypts per engineered intestine.

### Hydrogel stiffness measurement

Hydrogel stiffness was measured using an MCR 702e MultiDrive™ oscillatory shear rheometer (Anton Paar) in a parallel plate configuration with a PP25/P3 measuring plate. The hydrogel was prepared as described previously: TeloCol-6 Type I Collagen solution (5225-50mL, Advanced Biomatrix) was neutralized according to the manufacturer’s instructions and mixed with Matrigel (356231, Corning) at a 75:25 (v/v) ratio. Samples were immediately loaded at the center of a pre-cooled (10°C) Peltier plate, and the measuring geometry was lowered to a gap of 0.5mm. Storage modulus (G^*′*^) and loss modulus (G^*′′*^) were recorded during a temperature ramp from 10°C to 37°C. An oscillation frequency of 0.5Hz was used throughout, with a strain amplitude of 0.1% during heating and 0.2% after gel polymerization. The Young’s modulus (E) can be estimated from the storage modulus (G^*′*^), assuming the hydrogel is elastic and nearly incompressible, using the formula *E* = 2*G*(1 + *ν*) ≈ 3*G*, where Poisson’s ratio (*ν*) is approximately 0.5. Hydrogel measurements across multiple batches and replicates yielded a Young’s modulus of 700±150Pa.

### Statistical analysis

Statistical analysis was carried out with GraphPad Prism 10 (GraphPad software), following the details in the respective figure legend. The number of donors is indicated as ‘d=x’, whereas the number of independent engineered intestines is referred to as ‘n=x’ in the respective legend text. To check if the data followed a normal distribution, Shapiro-Wilk normality test was performed. When comparing two sets of data, unpaired two-tailed t-tests were carried out, including Welch’s correction when the SD determined not identical between the groups by a F test. In case the data was not normally distributed, Mann-Whitney sum-rank tests were performed. For comparisons involving more than two datasets and multiple variables, we performed one-way and two-way ANOVA, including Tukey’s multiple comparison or Dunnett’s comparison when comparing to a control condition. Data are visualized with their mean and standard deviation, where indicated, with the exact P values highlighting the statistical significance in the respective legend text.

### Single-cell dissociation of the engineered epithelium for scRNA-seq and scMultiome

Prior to dissociation, surfaces were cleaned with RNase AWAY™ Decontamination Reagent (10328011, Invitrogen) and all tubes were coated with a coating solution containing HBSS -Ca2+/ -Mg2+ (14175095, Gibco) + 1% BSA (130-091-376, Miltenyi Biotec) at 4°C for at least 15 min. Three independent engineered intestines were pooled for each scRNA-seq and five engineered intestines for scMultiome samples. Intestinal monolayers were detached from the matrix, collected and transferred to DNA LoBind 1.5mL tubes (22431021, Eppendorf) with 1 mL of coating solution using forceps. In the case of the scMultiome experiment, where the epithelium integrity was compromised, the epithelium and hydrogel matrix were apically punched using a 3 mm puncher and transferred to a 1.5 mL tube. The punched hydrogel was dissolved by incubating the mix with 1 mL of collagenase type I (100 U/mL, 17100017, Gibco) in Advanced DMEM/ F12 for 15 min at 37°C, following three washing steps in coating solution to clean the presence of collagenase in the sample and remove the digested hydrogel. Pooled and cleaned monolayers were pipetted up and down 50 times to mechanically disrupt the monolayers into smaller fragments. The single-cell dissociation was performed using the Neural Tissue Dissociation Kit (P) (Miltenyi Biotec, 130-092-628). Briefly, the samples were centrifuged at 400g for 5 min at 4°C. The supernatant was removed and the samples were incubated with 1 mL of the enzymeP Mix1 at 37°C for a total of 15 min, homogenizing the sample every 5 min by pipetting up and down a total of 50 times with a P1000 tip. After the first incubation, 30 µl of enzyme A Mix2 were added to the sample and incubated for 10 min, with homogenization steps every 5 min and inspection of the suspension under a bright field microscope. The single-cell suspension was filtered through a PluriStrainer Mini 40 µm (43-10040-40, Pluriselect), previously coated with coating solution, into new 1.5 mL DNA LoBind tubes and centrifuged at 450 g for 5 min at 4°C. The pellets were washed a total of three times with coating solution at 450 g for 5min at 4°C. Next, the samples were counted using the Countless 3 FL (AMQAF2000, Invitrogen) and adjusted to the desired concentrations for multiplexing, nuclei isolation or loading on the Chromium X (10x Genomics).

### Collection of TRMs for scRNA-seq

After collection of the epithelium using forceps, the remaining matrix and TRMs were punched using a 3 mm puncher and transferred to a 1.5 mL tube. The matrix was dissolved by incubating the sample with 1 mL of collagenase type I (100 U/mL, 17100017, Gibco) in Advanced DMEM/ F12 for 15 min at 37°C. After incubation, the sample was centrifuged at 500 g for 2 min at 4°C and washed with 1 mL of HBSS (-Ca^2+^/ -Mg^2+^) (14175095, Gibco) +1% BSA (130-091-376, Miltenyi Biotec). This step was repeated twice. Before final cell counting, Dead Cell Removal Kit (130-090-101, Miltenyi Biotec) was used following manufacturer instructions to remove apoptotic cells and lastly passed through a 40 µm Flowmi™ filter (BAH136800040, Millipore Sigma). Afterwards, the TRM cells were transferred to a 0.5 mL tube to facilitate visualization and accurate cell counting using the Countless 3 FL prior loading into the Chromium X (10x Genomics).

### Nuclei isolation for scMultiome

Nuclei isolation was performed based on 10x Genomics Demonstrated protocol CG000365 RevC. Briefly, single cells were centrifuged for 4 min at 400g and 4°C. The cell pellet was resuspended in cold lysis buffer (10 mM Tris-HCl pH7.4, 10 mM NaCl, 3 mM MgCl2, 1% BSA, 1 mM DTT, 0.1% Tween-20, 0.1% Nonidet P40 Substitute, 0.01% Digitonin, 1U/µl RNAse inhibitor). After 6 min on ice, cold wash buffer was added (10 mM Tris-HCl pH7.4, 10 mM NaCl, 3 mM MgCl2, 1% BSA, 1 mM DTT, 0.1% Tween-20, 1U/µl RNAse inhibitor) and nuclei were spun for 5 min at 500g and 4°C. The pellet was resuspended in cold wash buffer and nuclei were counted with Propidium Iodide (ViaStain AO/PI, Nexcelom Bioscience) using the Countess 3 FL (Thermo Fisher Scientific). After spinning, nuclei were resuspended in Diluted Nuclei Buffer (10x Genomics) at recommended concentrations for 10x Genomics Multiome. The quality of isolated nuclei was evaluated by microscopy with DAPI and bright-field.

### scRNA-seq and scMultiome library preparation

Singe-cell RNA-seq libraries were generated using 10x Genomics Chromium Next GEM Single Cell 3 v3.1 reagents according to user manual CG000315 Rev F. For multiplexed scRNA-seq, single-cell suspensions of 9 biological samples were individually tagged with 10x Genomics 3’ CellPlex Set A reagents according to the Demonstrated Protocol CG000391 Rev B Protocol 1. Equal amounts of tagged cells were pooled together for each sample. Three lanes of Chromium Next GEM Chip G were loaded targeting 30.000 cells per lane. Cell encapsulation and library preparation was performed according to user guide Chromium Next GEM Single Cell 3 Reagent Kits v3.1 with Feature Barcode technology for Cell Multiplexing CG000388 Rev C. Single-nuclei multiome was performed using 10x Genomics Chromium Next GEM Single Cell Multiome ATAC + Gene Expression reagents and corresponding user guide CG000338 Rev F. Two lanes were loaded per sample targeting 10.000 cells per lane. All single-cell libraries (scRNA, snRNA, Multiplex and snATAC) were sequenced on the Illumina NovaSeq 6000 platform.

### Read alignment of single cell genomic data

We ran Cell Ranger (v8.0.0) multi pipeline with 10x Genomics References 2024-A (hg38, GENCODE v44) as reference for read alignment, demultiplexing and gene quantification of multiplexed scRNA-seq data, Cell Ranger (v7.1.0) with 10x Genomics References 2024-A for TRM scRNA-seq data, Cell Ranger Arc (version 2.0.1) with 10x Genomics References 2020-A (hg38, GENCODE v32) for scMultiome data. We then loaded the Cell Ranger and Cell Ranger Arc pipelines filtered count matrix to Seurat (v4.3.0.1)^136^ and Signac (v1.10.0)^137^ for further preprocessing.

### Preprocessing and cell type annotation of scRNA-seq data

For the multiplexed scRNA-seq data of engineered epithelium, cells with fewer than 500 or more than 6,000 genes expressed, or total UMI counts exceeding 30,000, or mitochondrial gene percentage above 25% were removed. For the TRM data, we only retained cells with more than 500 genes, UMI counts between 1,000 and e10 and mitochondrial percentage less than 15%. For the epithelial data with and without TRM coculture, we only retained cells with more than 1,000 genes, UMI counts between 3,000 and 50,000 and mitochondrial percentage less than 25%.

For all datasets, raw counts were normalized using the ‘LogNormalize’ function with a scale factor of 10,000. 3,000 highly variable genes (HVGs) were identified using the variance-stabilizing transformation (‘vst’) method. Data were scaled and principal component analysis (PCA) was performed using these variable features. The first 20 principal components were used as input for ‘FindNeighbors’ and dimensional reduction with Uniform Manifold Approximation and Projection (UMAP). The resulting graph was used for ‘FindClusters’ with the Louvain algorithm at resolution 1.0.

For the multiplexed data, after inspection of regional marker gene expression, we excluded the potential foregut sample (“don55”) marked by SOX2 and a sample with mixed ileal and colonic identity (“IF17ile”, FABP6+ and SATB2+). We then removed ribosomal genes, mitochondrial genes, genes located on sex chromosomes and genes without proper gene symbols to avoid influence of these confounding factors to the integration of this dataset derived from multiple donors and reran the pipelines from normalization to clustering.

Epithelial cell type annotation was initialized on cluster level based on expression of canonical intestinal cell type and regional markers, including: stem cells (LGR5, OLFM4, SMOC2, ASCL2), absorptive cells (APOA4, FABP6, CEACAM7), goblet cells (MUC2, FCGBP), enteroendocrine cells (CHGA), transit amplifying cells (MKI67), BEST4+ cells (BEST4), epithelial regional identity markers (SATB2, GATA4, CDX2, PDX1) and leukocytes (PTPRC). Clusters with similar expression patterns were merged and resulted in a coarse-grained annotation.

For the multiplexed data which did not resolve a cluster representing distal small intestine stem cells, we then extracted stem cell clusters and absorptive cell clusters of all regions and refined the cell type classification along the continuum based on feature gene expression. We selected the top 10 genes with the highest correlation with summarized scaled expression of LGR5, SMOC2 and ASCL2 across stem cells and absorptive cells and calculated the stem cell score of each cell by summing the scaled expression of selected genes. We defined cells with stem cell scores higher than the 60th percentile of positive scores as stem cells and others as absorptive cells. To reduce the noise of classification solely based on scores of individual cells, we performed high-resolution clustering (resolution = 5), and defined cells belonging to the high resolution clusters with more stem cells than absorptive cells were defined as stem cells, regardless of their individual scores.

To resolve the goblet cell heterogeneity in the multiplexed data, we subsetted goblet cells from the entire dataset based on their coarse-grained cell type annotation. The subset was re-normalized, and variable features were recalculated and re-clustered (resolution = 1). We excluded clusters with enriched expression of MKI67 to get non-proliferative goblet cells and performed subclustering. We identified cluster markers using the ‘wilcoxauc’ function implemented by the ‘presto’ package and defined significant cluster markers with Wilcoxon rank sum test BH-adjusted P-value (‘padj’) < 0.05, expressed cell percentage difference (pct_in) > 10%, expression log fold change (logFC) > 0.1.

TRM cell type annotation was performed on the cluster level. We first annotated epithelial (EPCAM+), B cells (CD79B+), proliferative CD4+ T cells (MKI67+), then annotated non-proliferative T cell subtypes based on mapping to a public adult human intestine tissue immune cell reference with two complementary approaches. We identified cell type markers of the in vitro and tissue data separately using the ‘wilcoxauc’ function with cutoffs Wilcoxon rank sum test BH-adjusted P-value < 0.05, expressed cell percentage difference > 10%, expression log fold change > 0.1. We quantified the marker gene overlap significance between in vitro and tissue cell types using -log10 transformed Fisher’s two-sided exact test nominal P value. For each in vitro cell type, we got the top 3 tissue cell type with the strongest overlap, or the subset of top 3 if the significance difference between pairs of top 3 is larger than 10. In parallel, we calculated Spearman correlation between our in vitro and tissue cell types based on the markers defined in both systems. For each invitro cell type, we got the top 3 tissue cell type showing the highest transcription program similarity, or the subset of top 3 if the significance difference between pairs of top 3 is larger than 0.1. We then annotated the in vitro T cells as the overlap of overlap-based and similarity-based matched tissue cell types. Based on further inspection of individual canonical marker gene expression, we manually renamed “Activated CD4 T”, “Activated CD8 T” and “MAIT-like CD4” as “CD4 TRM”, “CD8 TRM” and “CD4 TRM Th17”.

### Construction of adult human intestine tissue transcriptome reference

To construct an adult human intestine tissue transcriptome reference, we collected the count matrix and cell type annotation of epithelial cells from three public adult intestine multi-region scRNA-seq datasets^35,138,139^. All datasets were processed using Seurat v4. After merging the datasets, we performed ‘LogNormalize’ normalization with a scale factor of 10,000. We identified HVGs independently for each dataset using the ‘vst’ method, selecting 3,000 HVGs per dataset. We then selected integration features across datasets using the ‘SelectIntegrationFeatures’ function. We ran principal component analysis (PCA) using the selected integration features and retrieved the top 50 PCs. We integrated the data with Cluster Similarity Spectrum (CSS)140 stratified by sample, and default setting for other parameters. We performed dimension reduction with UMAP, and Louvain clustering based on CSS embeddings using the ‘FindNeighbors’ and ‘FindClusters’ functions at a resolution of 1.0. We compared the data without batch effect correction, integration with Harmony or with CSS integration, and chose CSS-integrated results based on visual inspection of canonical cell type marker gene expression.

To establish a consistent cell type annotation scheme across datasets, we created a mapping between original annotations and standardized categories. We classified cells into major epithelial lineages: absorptive cells (enterocytes, colonocytes), goblet cells, BEST4+ cells, enteroendocrine cells (EECs), follicle-associated epithelium (FAE), stem cells, M cells, Paneth cells, secretory progenitors and tuft cells. We further refined these annotations by incorporating regional information. We defined the G2M-phased cells defined by Seurat ‘CellCycleScoring’ function within the clusters with enriched expression of MKI67 as the transit amplifying (TA) cells. Cell type identities were validated using established marker genes. For rare cell populations (e.g., Paneth cells and tuft cells), we calculated expression-based scores using canonical markers (DEFA5, DEFA6 for Paneth cells; RGS13, AVIL, ALOX5 for tuft cells) and assigned cells based on these scores. To be more specific, we identified the top 5 positively correlated genes with summarized DEFA5 and DEFA6 scaled expression profiles across epithelium among the 3,000 HVGs as feature genes of Paneth cells. We summed up the scaled expression levels of the selected top 5 Paneth-associated genes for each cell as the Paneth cell feature score, and defined cells with scores exceeding 97.5th percentile of all positive scores as Paneth cells. We applied a similar procedure to identify Tuft cells, with 10 Tuft cell feature genes and using 95 percentile as the selection cutoff.

To establish a tissue reference for colonocyte subtype annotation in the Salmonella dataset, we subsetted to non-small intestine TA cells, stem cells and colonocytes, integrated the data with CSS and performed subclustering based on the CSS embeddings at resolution = 0.4. We identified cluster markers using the ‘wilcoxauc’ function in the ‘presto’ package, with cutoffs padj < 0.05, pct_in > 10, logFC > 0.1. We selected the top 100 markers ranked by logFC from each cluster and unified them as features for comparison between in vitro and tissue colonic stem cells and colonocytes.

### Preprocessing and cell type annotation of scMultiome data

To create a unified peak list across samples, we called open chromatin accessible regions in each sample using MACS2 (v2.2.9.1) with ‘CallPeaks’ function and then quantified them with ‘FeatureMatirx’ in ‘Signac’. We only keep regions located on standard chromosomes. We annotated peaks to most nearby genes using the ‘annotatePeak’ function in ‘ChIPseeker’ based on Ensemble (v98) annotation. We only retained cells with ATAC UMI count between 200 and 50,000, RNA UMI count less than 20,000, RNA gene number between 500 and 5,000, mitochondrial percentage less than 50%, nucleosome signal (calculated by ‘NucleosomeSignal’) less than 1, transcription start site (TSS) enrichment score (calculated by ‘TSSEnrichment’) less than 1 and hg38 genome blacklist ratio less than 0.5%. We ran the same pipeline of scRNA-seq data from read count normalization to dimension reduction and clustering as described in the previous section (“Preprocessing and cell type annotation of scRNA-seq data”). For the scATAC-seq data, we performed term frequency-inverse document frequency (tf-idf) normalization on read counts using the ‘RunTFIDF’ function. We defined the regions with over 1% detection rate as top features and ‘RunSVD’ based on the top features. We projected cells into two dimension UMAP embeddings and ‘FindNeighbors’ based on top 2 to 50 latent semantic indexing (LSI).

We performed major epithelial cell type annotation on RNA cluster level based on canonical cell type marker expression. We subsetted to colonic stem cell and colonocyte clusters, calculated the Spearman correlation coefficients between in vitro and tissue stem cell to colonocytes clusters using the in vitro expressed colonic stem cells and colonocytes feature genes defined in the tissue reference. We performed hierarchical clustering on the in vitro clusters based on the similarity profiles and classified them into stem cells and four colonocyte subtypes indicating differential maturity accordingly. Colonocyte clusters with mostly exclusive presence in the Salmonella infected sample were defined as infection-specific colonocyte clusters. We transferred RNA level cell type annotations to ATAC data, and merged subtypes that could not be clearly resolved by ATAC data as annotations on the ATAC level.

### Identification of differential features between conditions

We performed differential analysis for co-clustered *S*.Tm infection and control cells in each cluster of stem cells, colonocytes and goblet cells, which have sufficient cell numbers for robust estimation. For infection-specific clusters which have less than 20 control cells in the clusters, we first identified matched control sample cells for each infected sample cell using the ‘get.knnx’ of the ‘FNN’ package with ‘k=1’ in the top 20 PC space. We identified differentially expressed genes between matched control and infected cells using the ‘wilcoxauc’ function by ‘presto’ package, with cutoffs padj < 0.05, pct_in > 10, logFC > 0.1. We applied the same procedure to identify differentially expressed genes in epithelium induced by TRM coculture except that we don’t need to use the kNN search approach to get matched control cells since there is no condition-specific cluster.

To identify differentially accessible regions (DARs) between Salmonella-infected and control conditions, we implemented a statistical framework based on linear regression modeling of binarized chromatin accessibility data, with sequencing depth (UMI counts) as covariate. Specifically, for each open chromatin region *i*, we fitted:

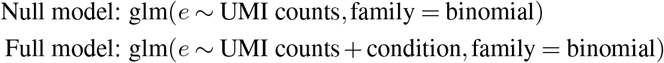

where *e* represents the binarized accessibility state vector for region *i* across all cells. We applied a Chi-square test comparing the two models using ANOVA. Regions are considered differentially accessible if they have Benjamini-Hochberg adjusted p-values below 0.05 and show at least a 0.5% difference in the proportion of cells with the region accessible between Salmonella-infected and control conditions.

### GRN construction and TFBS motif enrichment

We reconstructed the gene regulatory networks from the scMultiome data along the stem-cell-to-coloncyte trajectory and each of the stem cell, colonocyte and goblet cell types shared by both control and infection conditions using Pando^141^ separately. We used the infection induced differentially expressed genes and 5,000 HVGs of each data as input HVGs for Pando. We constrained our candidate regulatory regions to be either conserved in mammals (defined in ‘phastConsElements20Mammals.UCSC.hg38’ provided by Pando) or overlap with ENCODE regions (defined in ‘SCREEN.ccRE.UCSC.hg38’ provided by Pando), and showing significantly correlated profiles with HVGs as output by ‘LinkPeaks’ in ‘Signac’ at the default distance threshold 500kb. We ran ‘find_motifs’ and used the Pando-provided transcription factor binding site (TFBS) motif database compiled from JASPAR and CIS-BP as reference for Pando to identify TFBS in candidate regulatory regions. We ran the ‘infer_grn’ function with a gaussian generalized linear model (model=‘glm’) to infer regulatory coefficients from log-normalized transcript counts and binarized peak counts summarized to SLM algorithm defined ATAC clusters (resolution = 10). We identified TF-target gene modules using the ‘find_modules’ function with default settings. We combined the modules from different datasets as inferred TF-target pairs.

For visualization, we selected the condition-dependent DEGs with the top 500 absolute value of expression log-transformed fold change between conditions and the union set of top 3 correlated Pando predicted TFs of each selected DEGs. We calculated pairwise Pearson correlation coefficients of expression profiles across cell types and conditions between selected genes, reduced the dimension of obtained correlation matrix to 20 principle components using ‘prcomp_irlba’ in the ‘irlba’ package and further projected genes to two dimension embeddings using ‘umap’ in the ‘uwot’ package with the obtained PCs as input.

We ran ‘AddMotifs’ function in ‘Signac’ to add the TFBS motifs information on our detected open chromatin regions, and obtained a motif occurrence matrix. To identify TFBS motifs over-represented in open chromatin regions with increased accessibility upon infection in each cell type, we ran ‘FindMotifs’ in ‘Signac’ on top 200 regions upregulated accessible regions, with 30,000 open chromatin regions with matched overall GC content retrieved by ‘MatchRegionStats’ function as background.

### CellChat analysis of epithelial-immune cell interactions

We used CellChat (v2)^142^ to infer intracellular communications between cocultured intestinal epithelial cells and tissue-resident memory T cells (TRMs). Normalized expression data was extracted from the epithelium and TRM combined Seurat RNA assay, and cell type labels were used to group cells for interaction analysis. We used the built-in human CellChatDB database as reference for ligand-receptor interactions. To reduce computational cost, we subsetted the expression data to include only signaling genes present in the database. Overexpressed genes and interactions were identified using the ‘identifyOverExpressedGenes’ and ‘identifyOverExpressedInteractions’ functions. Communication probabilities were computed using the ‘computeCommunProb’ function with ‘type = truncatedMean’ and ‘trim = 0.05’. We subsetted to interactions between proximal small intestinal enterocytes and immune cell types, excluding proliferating CD4+ T cells for visualization.

**Supplementary Table 1**. Key resource table.

**Supplementary Table 2**. Single cell sequencing data quality of cells passed filterings.

**Supplementary Table 3**. Differential features between cell types and conditions, including a) Top 100 cell type marker genes of engineered intestinal epithelium; b) Top 500 enterocytes (duodenum, ileum) and colonocytes (colon) marker genes; c) Top 100 marker genes of non-proliferative goblet cell clusters; d) Top 100 cell type marker genes of epithelium in control and *S*.Tm condition combined data; e) Differentially expressed genes (DEGs) in epithelium upon *S*.Tm infection; f) Co-expression gene modules of DEGs upon *S*.Tm infection; g) Differentially expressed genes in epithelium upon TRM coculture.

**Supplementary Table 4**. Functional enrichment results, including a) Biological process (BP) and Molecular function (MF) terms with enrichment scores of enterocyte and colonocyte marker genes; b) BF enrichment of non-proliferative goblet cell cluster markers; c) Nominal hypergeometric test P values of KEGG pathway and MSigDB Hallmark gene set enrichment of per cell type DEGs of epithelium upon infection; d) Nominal hypergeometric test P values of KEGG pathway enrichment of infection-induced DEG co-expression gene modules; e) DAVID output of epithelial up-regulated genes upon TRM coculture.

**Supplementary Table 5**. Protein ranks of colonic luminal content, including a) Identified proteins in colonic luminal content from three donors, with abundance, IBAQ and rIBAQ values. b) Averaged protein ranks based on averaged IBAQ values across donors.

**Supplementary Table 6**. Transcription factor binding site motif enrichment in top 200 open chromatin regions with increased accessibility in infection-specific colonocyte subtype 2 upon *S*.Tm infection.

**Supplementary Table 7**. Interaction probability of cell type enriched ligand-receptor pairs inferred by CellChat.

**Supplementary Video 1**. TRMs migration (pink) in the top-layer of the duodenal epithelial monolayer (blue).

**Supplementary Video 2**. TRMs flossing (pink) in the top-layer of the duodenal mScarlet3-CDH1+ epithelium (green) around crypt invaginations and along the crypt-villi axis.

**Supplementary Video 3**. 3D-rendered video capturing the interaction between TRM (yellow) with duodenal mScarlet3-CDH1+ epithelial cells (red) in the engineered tissue.

**Extended Data Figure 1.**
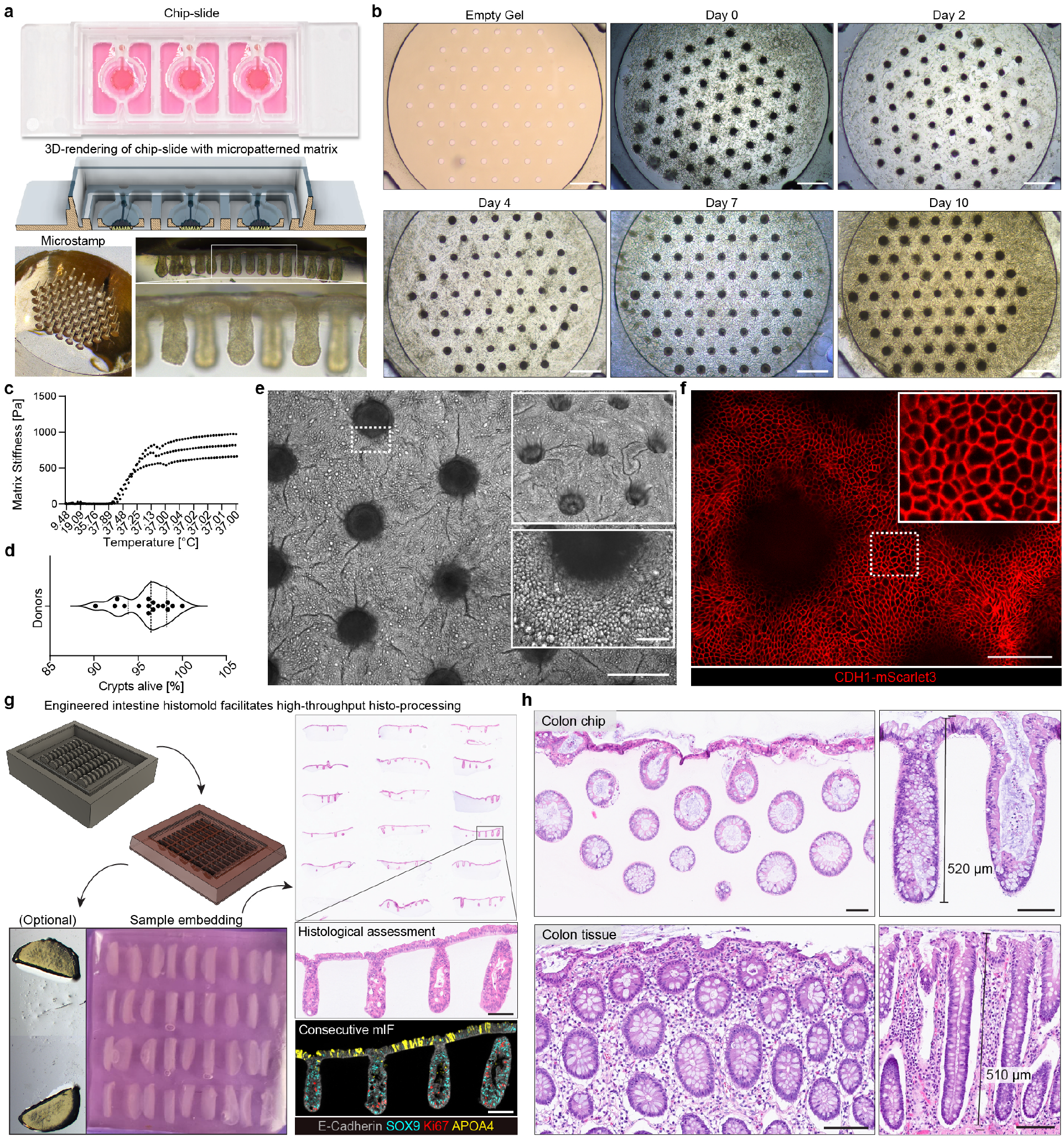
Generation and assessment of engineered intestinal epithelium using high-throughput histology. a) Picture of a chip-slide with culture medium, including a 3D-rendered image of the cross-section of the chip-slide containing three individual chip segments and an image of the micropatterned stamp with a representative bright-field image of an engineered intestine cross-section. b) Time course of biomimetic epithelium formation from the stamped empty ECM to epithelium coverage and maturation. Scale bars: 500 µm. c) Rheological characterization of the collagen-Matrigel mixture used as the ECM in the engineered intestines. Each dotted-line represents a technical replicate of the hydrogel-mixture. d) Quantification of viable crypts of 16 donors from duodenum, ileum and colon. e) Representative EDF brightfield image of a Z-stack, showing the top-view of a day-9 engineered epithelium. Scale bar: 200 µm. The magnified insert (top) shows a 3D-reconstruction of the engineered epithelium with crypts penetrating into the matrix. The magnified insert (bottom) shows an EDF image of a Z-stack in the crypt area. Scale bar: 50 µm. f) Representative EDF fluorescent image of selected Z-stacks with magnified insert, showing an iCas9 CDH1-mScarlet3 day-9 duodenal engineered epithelium with tight E-Cadherin+ adherens junctions (red). Scale bar: 100 µm. g) Overview of the high-throughput histo-processing workflow using 3D-printed histomold facilitating co-planar embedding of up to fifty engineered epithelia at once. Post-FFPE embedding, dozens of engineered epithelia can be visualized within one section as illustrated in a representative H&E. Consecutive sections subjected to H&E and mIF to evaluate the spatial organization and maturation of a duodenal engineered epithelium at day 6. Scale bar: 100 µm. h) H&E cross-sections of representative colon engineered epithelium and primary tissue, showing high morphological similarity, crypt depth, and goblet cell distribution. Scale bars: 100 µm.

**Extended Data Figure 2.**
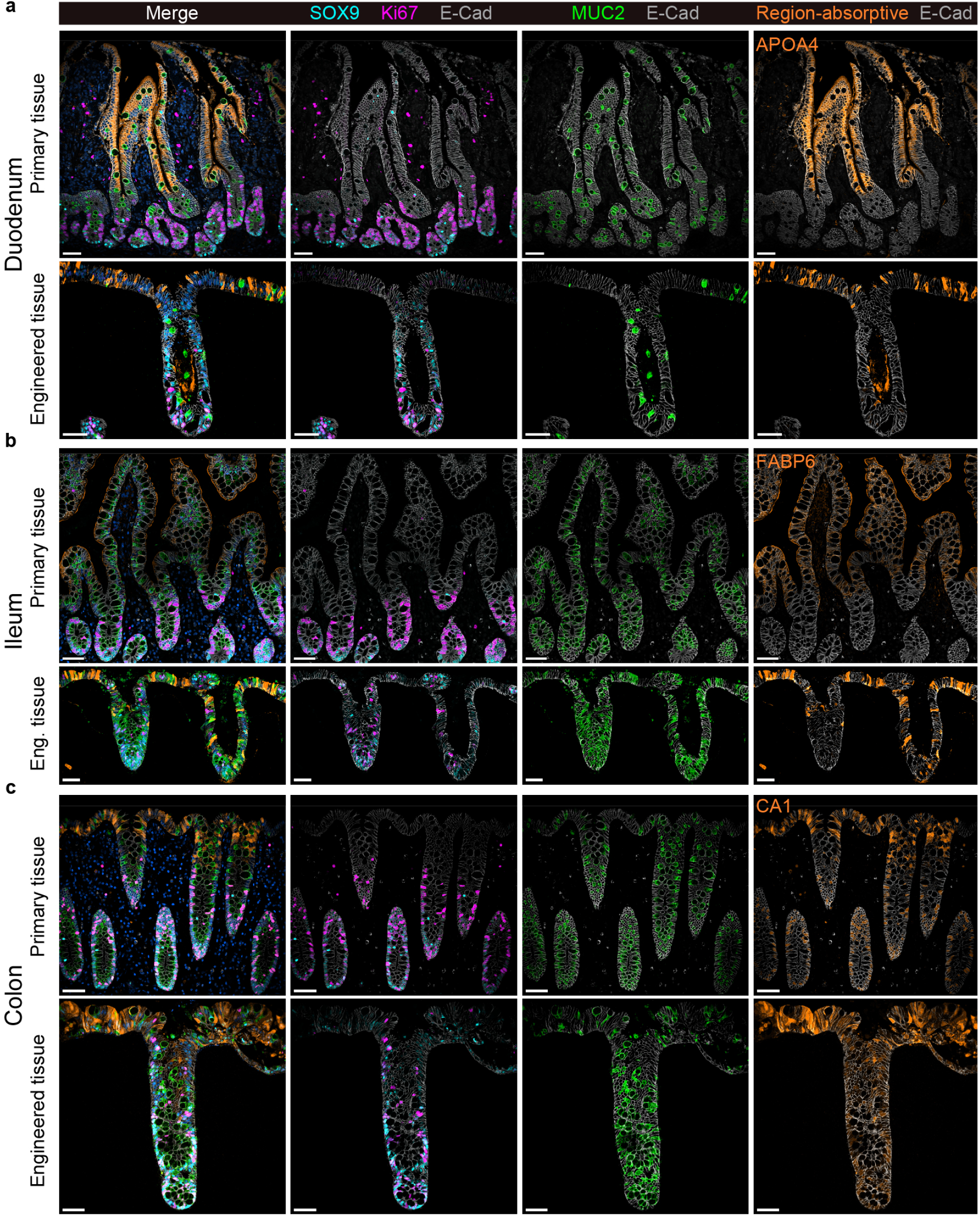
Region-specific patterned differentiation recapitulates physiological maturation of the organ of origin. a-c) Representative mIF cross-section of engineered duodenal, ileal and colonic epithelia in direct comparison to the donor-matched primary tissue (ileum not matched), highlighting the stem-niche specific SOX9+ (cyan) and transit-amplifying Ki67+ cells (pink), matching abundance of MUC2+ goblet cells (green) and high expression of the region-enriched absorptive lineage marker (orange) in the E-Cadherin+ epithelium (grey). The duodenal epithelium was marked by strong expression APOA4+ enterocytes, matching the parental tissue. Expression of distal FABP6+ enterocytes in the top-layer of the engineered epithelium matched ileal tissue. The colonic epithelium exhibited CA1+ colonocytes at comparable levels to the organ of origin. Scale bars: 50 µm.

**Extended Data Figure 3.**
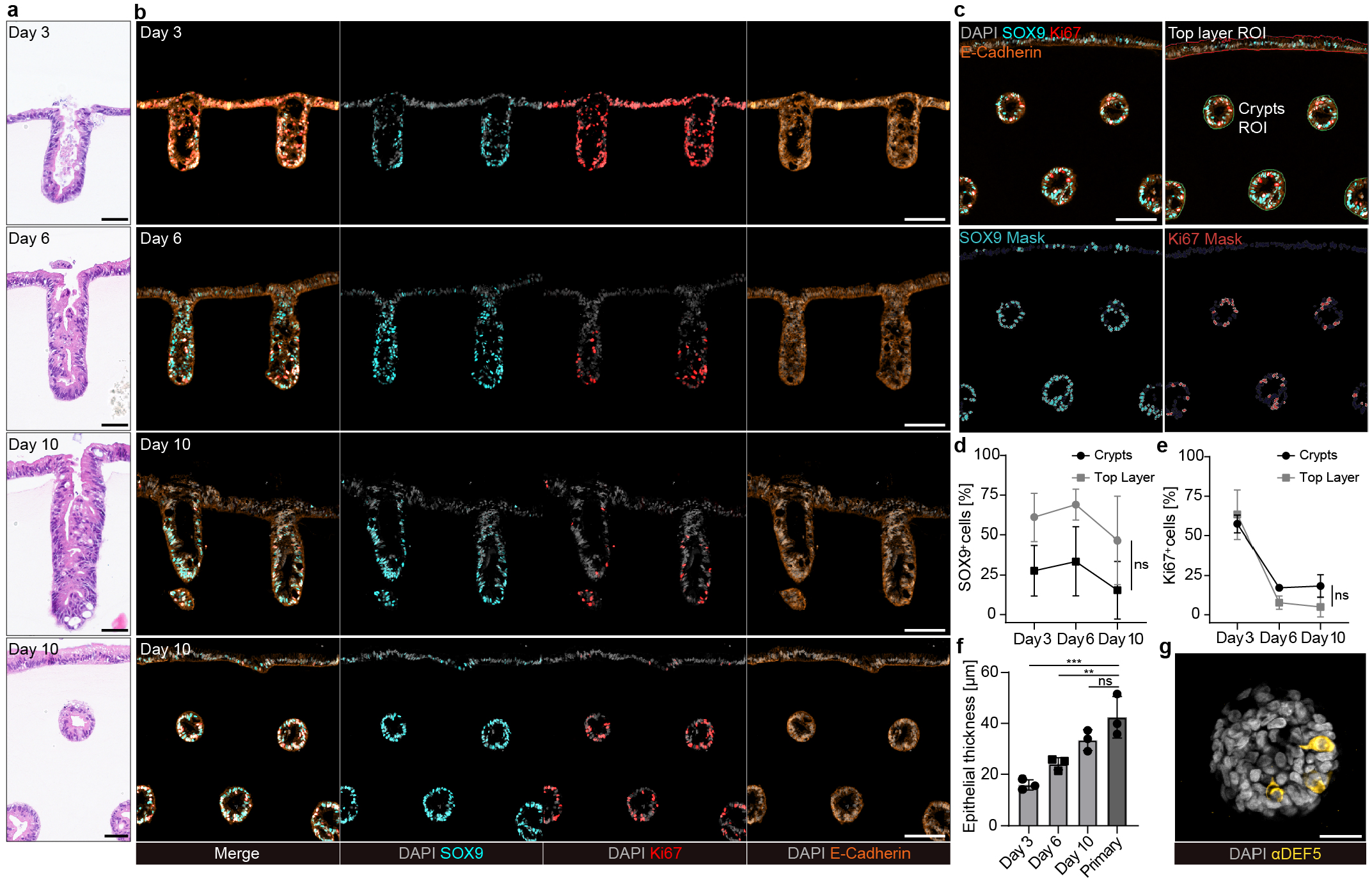
Differentiation assessment over time reveals robust spatiotemporal patterning in engineered intestines. a) Representative H&E images of engineered duodenal intestines over 10 days of maturation, highlighting epithelial polarization, thickening, and increasing crypt depth. Scale bar: 50 µm. b) Representative mIF images of spatiotemporal patterning of SOX9+ (cyan) and Ki67+ (red) cells of E-Cadherin+ (orange) duodenal epithelium over 10 days. Scale bar: 100 µm. c) Representative mIF showcasing crypt and top layer annotations and cell segmentation with respective masks for SOX9+ (cyan) and Ki67+ cells (red). Scale bar: 100 µm. d) Quantification of SOX9+ cells in the respective annotation across multiple time points, normalized to overall DAPI+ cell count in the annotation. e) Quantification of Ki67+ cells in the respective annotation over time, normalized to overall DAPI+ cell count in the annotation. Line plots in d-e) represent the mean of donors (d=3) +/-SD at day 9, derived from three engineered intestines per donor. Two-way ANOVA with Tukey’s multiple comparisons test (ns=P>0.05). f) Average epithelial thickness in donor-matched duodenal tissue and engineered epithelia. Bars represent the mean of donors (d=3) +/-SD (dots), derived from regions of engineered epithelia (n=3) per donor and matched primary tissue (dots). Non-paired one-way ANOVA with Dunnett’s multiple comparisons test compared to the primary tissue as control, non-significant (ns=P>0.05, **P=0.0044, ***P=0.0004). g) Whole mount staining of the crypt bottom of a duodenal engineered epithelium at day 20, with DEFA5+ Paneth cells. EDF fluorescent image of a Z-stack. Scale bar: 25 µm.

**Extended Data Figure 4.**
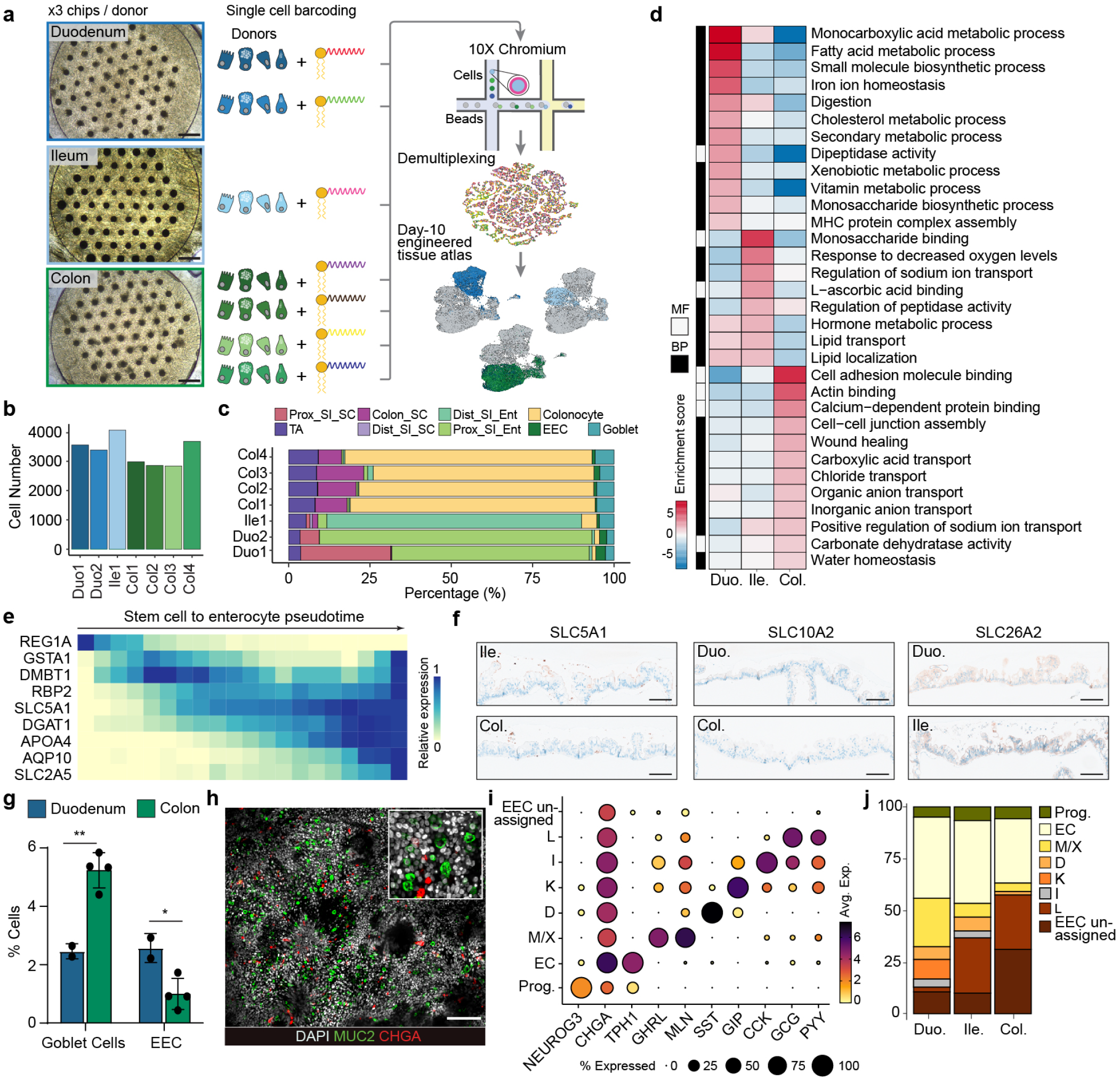
scRNAseq of engineered intestinal epithelium reveals high reproducibility and region-specific cell distributions and functions. a) Schematic of the multiplexed scRNA-seq characterization of day-10 engineered intestinal epithelia from duodenum, ileum and colon. b) Recovered cells from each of the sequenced donors, highlighting the accuracy of the dissociation and equal contribution of each donor to the resulting atlas. c) Stacked barplot showing cell-type percentages of each of the individual donors, underscoring the reproducibility of the model. d) Heatmap showing a selected set of enriched biological processes (BP) and molecular functions (MF) derived from the top 500 marker genes of enterocytes (duodenum, ileum) and colonocytes (colon). The enrichment score for each term is calculated by subtracting the significant -log10 transformed hypergeometric test BH-corrected P value of the region of interest and the average of those from the other two regions. e) Heatmap showing expression patterns of adult duodenum tissue zonation markers along stem cell to enterocyte diffusion map inferred trajectory of the engineered epithelium. f) Complementary to Fig. 1l. IHC stainings of SLC5A1, SLC10A2, and SLC26A2 in one cross-section containing duodenal, ileal and colonic engineered epithelia. Scale bar: 100 µm. g) Comparison of enteroendocrine cell (EEC) and goblet cell proportions on the engineered epithelia, derived from the day-10 scRNA-seq dataset and coloured by the region of origin. Bars represent the mean +/-SD. dDuo=2 and d=4Col (dots), n=3 pooled per donor prior sequencing, unpaired two-tailed t test (*PEEC=0.0262, **PGC=0.0039). h) Whole mount staining image showing goblet cells (green) and EECs (red) on the engineered epithelium. EDF fluorescent image of a Z-stack. Scale bar: 100 µm. i-j) (i) Dot plot showing the average expression of the main intestinal EEC-derived hormones at day 10. The expression is shown for the subclustered EEC subpopulations, identified as different subtypes. (j) Stacked barplot showing the proportions of the subclustered EEC populations within their region of origin.

**Extended Data Figure 5.**
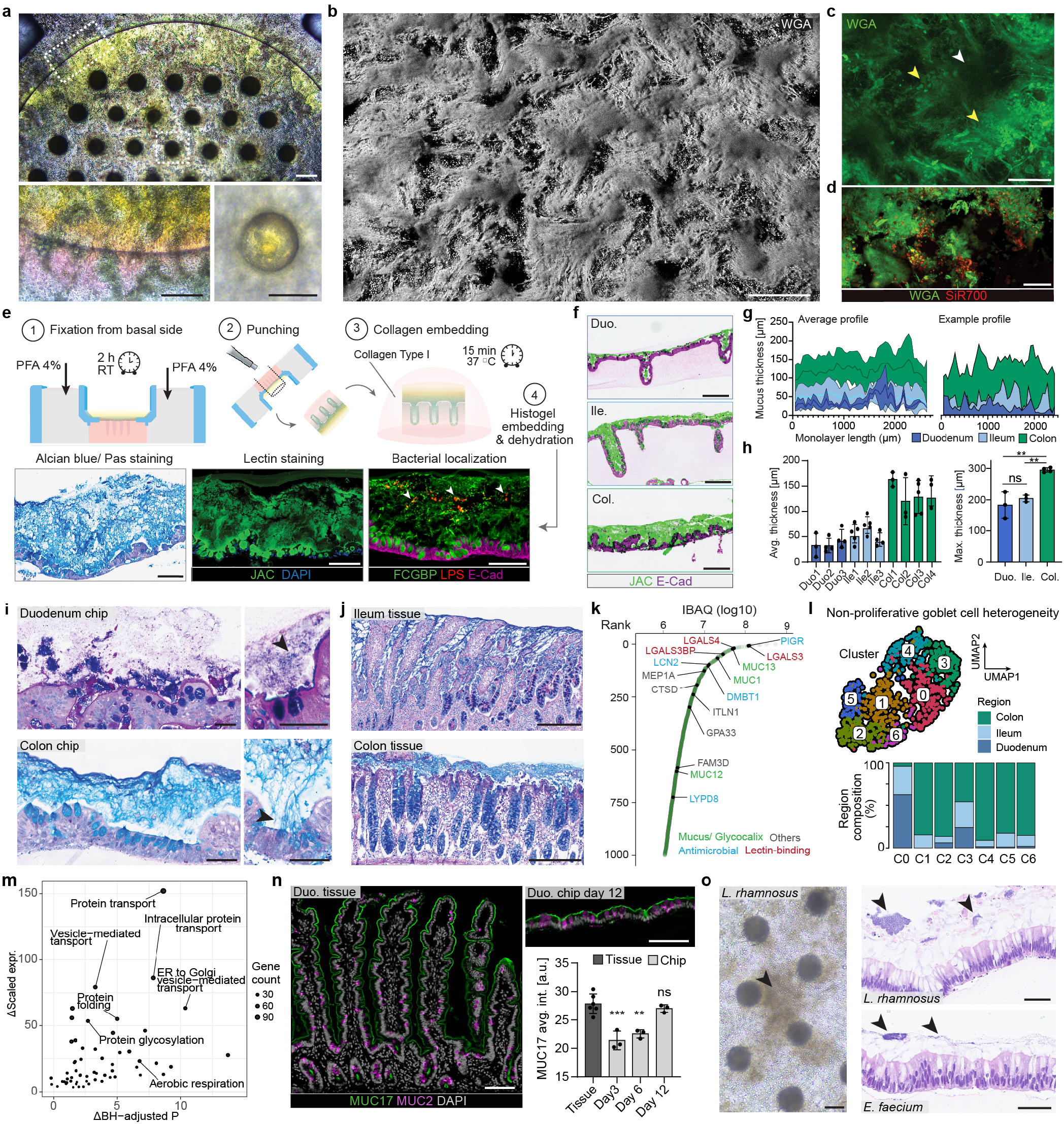
Characterization and functional validation of a mucus layer with in vivo-like region-specific properties. a) Representative bright field image of a day-9 colonic engineered epithelium with accumulated mucus (4 days, 5 µl; yellow; top). Magnified inserts show the side of the epithelium and one crypt, where mucus accumulates densely acquiring a yellow hue (yellow; bottom). Scale bars: 200 µm (top, bottom left), 100 µm (bottom right). b-c) Fluorescent live image showing a 3D reconstruction from a Z-stack of a day-13 ileal engineered epithelium with accumulated mucus (5 days, 5 µl), stained with Wheat Germ Agglutinin (WGA; grey). Scale bar: 200 µm (b). Maximum intensity projection image of a Z-stack derived from (c), showing WGA-stained mucus (green) and shed cells (yellow arrows) at the crypt opening (white arrow). Scale bar: 100 µm (c). d) Fluorescent live image of luminal content showing mucus (green) with shed cells (red). Scale bar: 100 µm. e) Schematic of the mucus preservation steps prior to histo-processing and FFPE preparation. A collagen layer on top of the punched epithelium facilitates preservation of the luminal content prior to histogel embedding. The method allows to study spatial characteristics of the mucus and luminal bacteria. Scale bars: 100 µm. f) Representative mIF of cross-sections of duodenal, ileal and colonic engineered epithelium with accumulated mucus (5 days, 30 µl) using the previously mentioned workflow from (e). Scale bars: 250 µm. g) (left) Average mucus layer thickness profile along common margins of cross-sections of day-9 duodenal, ileal and colonic engineered epithelia with accumulated mucus (5 days, 30 µl). Data obtained from d=3-4 and n=3-5 measured per donor and pooled together within each region. Each engineered intestine (n) was measured and averaged in two different cross-sections. Line connects average thickness of each location +/-SD represented by shaded area. (right) Example mucus layer thickness profile along common margins of one engineered epithelium per region. h) (left) Quantifications of the average mucus layer thickness in day-9 duodenal, ileal, and colonic engineered epithelia with accumulated mucus (5 days, 30 µl) from different donors. Bars represent the mean +/-SD of n=3-5 per donor measured and averaged in two different cross-sections (dots). (right) Maximum thickness measured point of the mucus layer per region. Bars represent the mean +/-SD of d=3-4 per region (dots) with n=3-5 measured per donor in two different cross-sections. The maximum measured point per donor corresponds to the maximum from the n=3-5 measured. One-way ANOVA with Tukey’s multiple comparison test (**P_Col.-Ile._=0.0039, **P_Col.-Duo._=0.0012, ns=P>0.05) (right). i-j) Representative AlcianBlue/ PAS staining of duodenal and colonic engineered epithelia with accumulated mucus (i) and ileum and colon primary tissues (j). The images show differences between regions in the structure of the mucus layer, the AlcianBlue/PAS+ ratio, and goblet cell mucus secretion (arrows). Scale bar: 50 µm (left), 250 µm (right). k) Ranked abundances of proteins detected in the luminal content of colonic engineered epithelia with accumulated mucus (5 days, 5 µl). Proteins that belong to the mucus proteome are highlighted by categories. l) UMAP of the non-proliferative goblet cells (-MKI67+) from the day-10 engineered tissue scRNAseq atlas, showing six different clusters (top) with different regional proportion contribution (bottom). m) Scatter plot showing the Gene Ontology (GO) (X-axis) and expression enrichment (Y-axis) of C1 marker enriched GO terms. Gene ontology enrichment was calculated as the difference between -log10 transformed hypergeometric test BH-corrected P value of C1 cluster markers versus average of those of other clusters. Expression enrichment was calculated as the difference between summarized scaled expression of GO term associated with C1 cluster genes versus the average of other clusters. n) Representative mIF staining of a duodenal tissue (left) and day-12 engineered duodenal epithelium (top right) cross-section, showing a comparable glycocalyx (green). Average MUC17 glycocalyx signal intensity (bottom right) in the duodenal tissue and engineered epithelia. Bars represent the mean +/-SD. d= 1 donor, n= 5 tissue regions from the same section and 3 engineered epithelia (dots), non-paired one-way ANOVA with Tukey’s multiple comparison test (compared to tissue: ***P_Day3_=0.0003, **P_Day6_=0.0013, ns=P>0.05). Scale bars: 100 µm. o) Representative bright field image of a Lacticaseibacillus rhamnosus biofilm on top of ileal engineered epithelia after 24 h of co-culture (left). Representative H&Es of ileal engineered epithelia with accumulated mucus (4 days, 30 µl) with visible Lacticaseibacillus rhamnosus and Enterococcus faecium biofilms after 24 h of co-culture (arrows; right). Scale bar: 100 µm (top), 50 µm (bottom).

**Extended Data Figure 6.**
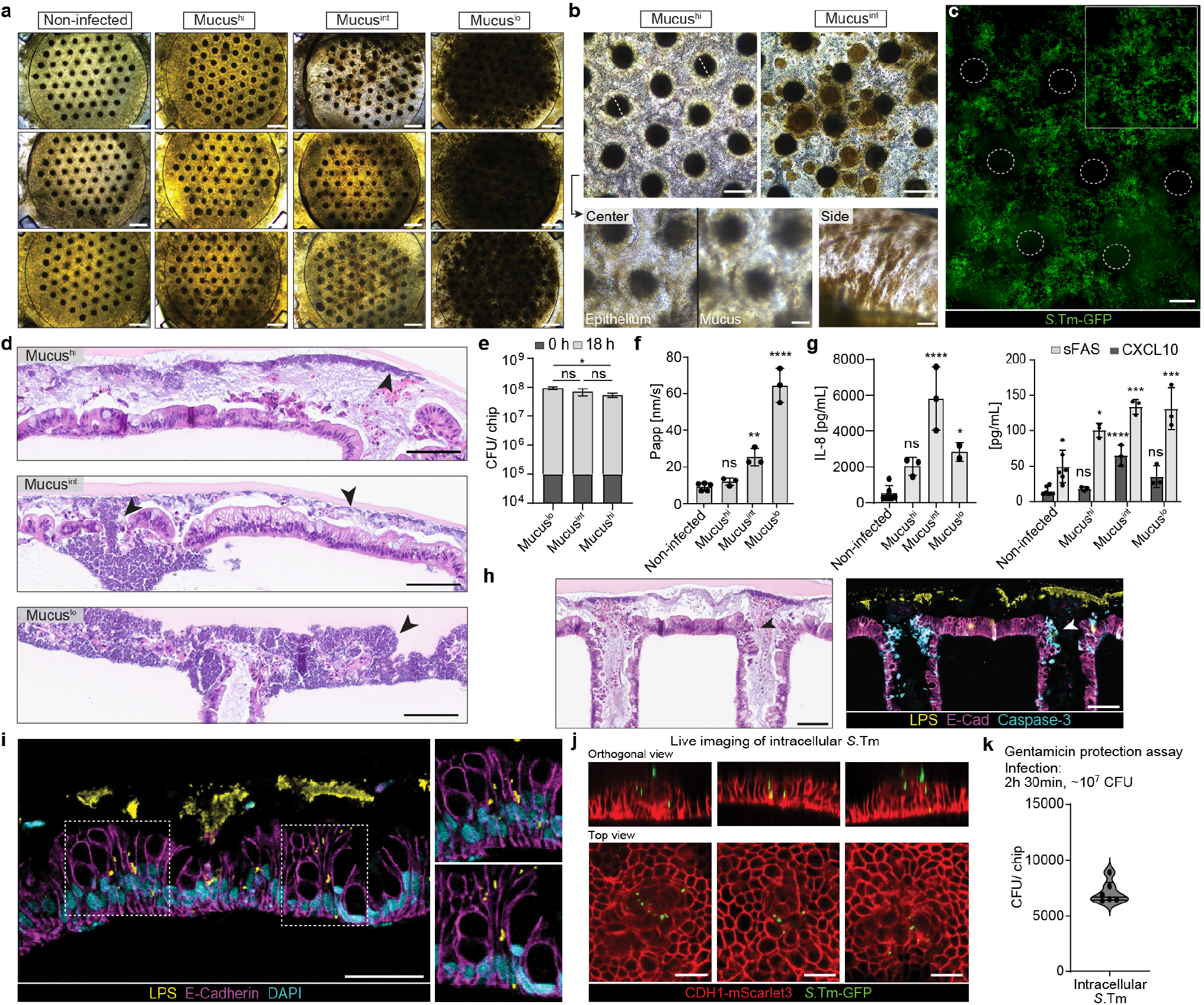
The colonic mucus layer influences infection dynamics and protects against *S*.Tm. a) Bright field images of non-infected and *S*.Tm-infected day-10 colonic engineered epithelia with different levels of accumulated mucus after 18 h of co-culture. Mucus accumulation: Mucus^hi^ (4 days, 5 µl), Mucus^int^ (2 days, 5 µl), and Mucus^lo^ (4 days, 120 µl, +wash). Scale bars: 500 µm. b) Representative bright field images of Mucus^hi^ and Mucus^int^ conditions after 18 h of *S*.Tm infection (top). Magnified areas of the Mucus^hi^ epithelium, observing the presence of bacteria in different planes with respect to the epithelium in the center and on the sides of the engineered intestines (bottom). Scale bars: 200 µm (top), 100 µm (bottom). c) Representative EDF fluorescent image of a Z-stack, showing a *S*.Tm biofilm (green) co-cultured with an epithelium with accumulated mucus (4 days, 5 µl). Scale bar: 100 µm. d) Representative H&E images of colonic Mucus^hi^, Mucus^int^, and Mucus^lo^ conditions showing a degree of epithelial damage by *S*.Tm (dark blue, arrows) that inversely correlates with the accumulated mucus level. Scale bars: 100 µm. e) *S*.Tm colony forming unit (CFU) counts from Mucus^hi^, Mucus^int^, and Mucus^lo^ samples at 0 h and 18 h of infection. Bars represent the mean +/-SD. d=1, n=3, non-paired one-way ANOVA with Tukey’s multiple comparison test (*P_MucusHI-MucusLO_=0.026, ns=P>0.05). f) Apparent permeability of non-infected, Mucus^hi^, Mucus^int^, and Mucus^lo^ samples after 18 h of *S*.Tm infection. Bars represent the mean +/-SD. d=1, n=3-5 (dots), non-paired one-way ANOVA with Dunnett’s multiple comparison test (compared to non-infected:****P_MucusLO_<0.0001,**P_MucusINT_=0.0031, ns=P>0.05). g) IL-8, CXCL10, and sFAS cytokine measurements from basal supernatants of non-infected, Mucus^hi^, Mucus^int^, and Mucus^lo^ samples after 18 h of *S*.Tm infection. d= 1, n=2-5, non-paired one-way ANOVA with Dunnett’s multiple comparison test (compared to non-infected: ****P_MucusINT-IL8_<0.0001, *P_MucusLO-IL8_=0.0272, ***P_MucusLO-sFAS_=0.0006, ***P_MucusINT-sFAS_=0.0004, *P_MucusHI-sFAS_=0.0164, ****P_MucusINT-CXCL10_<0.0001, ns=P>0.05). h) (left) Representative consecutive H&E and mIF stainings of a Mucus^hi^ sample, showing shed (black arrow) and CASP3+ (white arrow) cells that accumulate in the crypt plume, blocking *S*.Tm entrance (dark blue). Scale bar: 100 µm. i) mIF image of a Mucus^int^ sample, showing areas of the epithelium with intracellular *S*.Tm and luminal biofilms. Scale bar: 50 µm. j-k) EDF fluorescent images of selected Z-stacks, showing an iCas9 CDH1-mScarlet3 duodenal reporter-line (red) infected with *S*.Tm (green) for 2 h 30 min prior gentamicin wash. Regions of the epithelium show intracellular *S*.Tm after 18 h post-gentamicin addition (left; scale bar: 20 µm). CFU counts of intracellular *S*.Tm after 18 h post-gentamicin wash. d=1, n=6 (right).

**Extended Data Figure 7.**
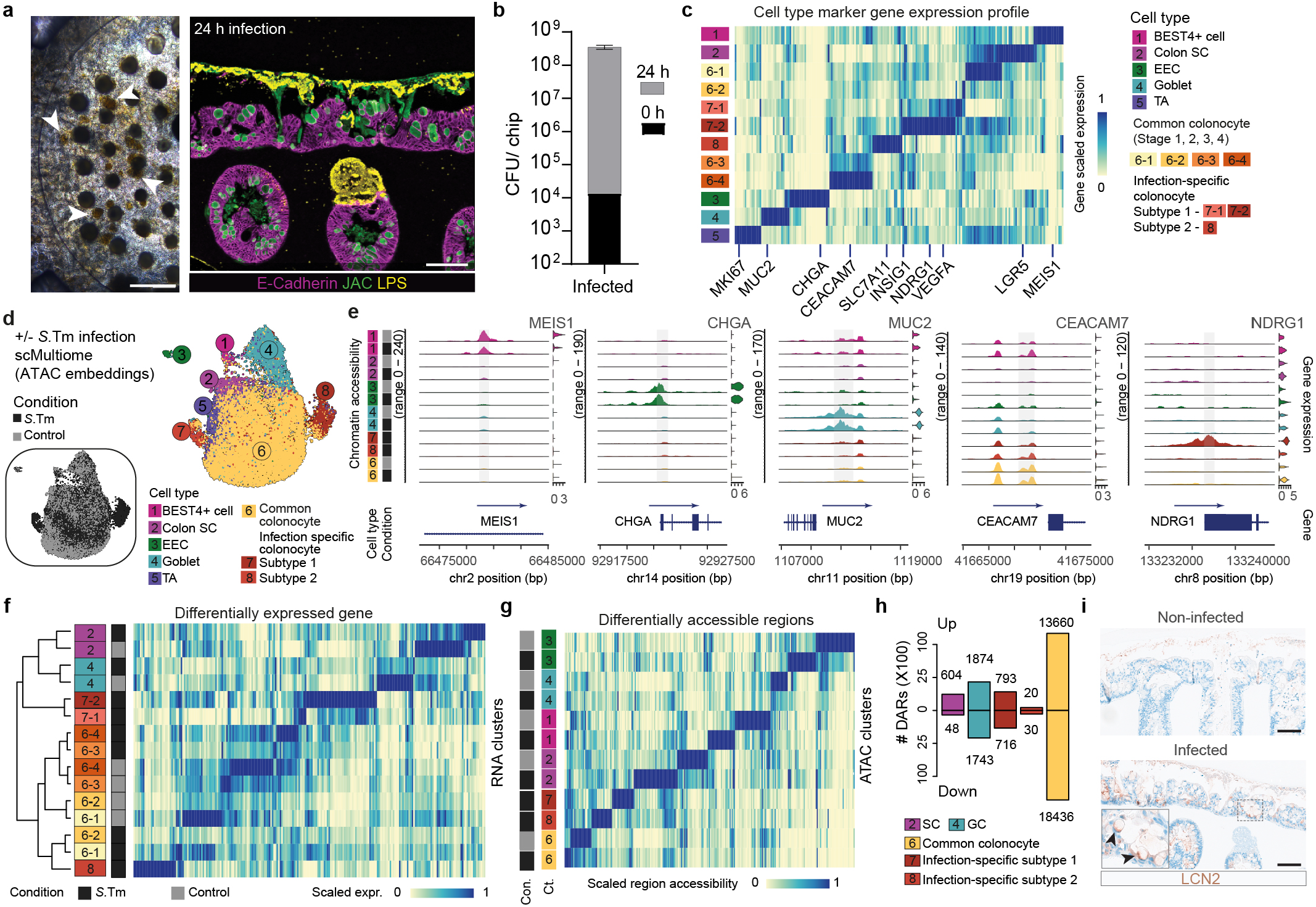
General colonic epithelium response to *S*.Tm infection. a) Representative bright field and mIF image of engineered colonic monolayers with accumulated mucus (4 days, 5 µl) after 24 h of *S*.Tm (yellow, LPS) infection, prior dissociation and nuclei isolation. White arrows indicate *S*.Tm colonies attached to the epithelium which caused barrier breach. Scale bars: 500 µm (left), 100 µm (right). b) CFU counting of *S*.Tm at 0 h and 24 h of infection, prior single-cell dissociation and nuclei isolation. Bars represent the mean +/-SD. d=1, n=3. c) Heatmap showing the expression patterns of marker genes across RNA clusters. d) UMAP of scATAC-seq data of colonic epithelium in control and *S*.Tm conditions, with cells colored by condition (left) or cell type. e) Violin plots showing cell type marker gene expression and coverage plots showing normalized chromatin accessibility of their promoter or nearby regulatory regions across cell types of different conditions: BEST4+ cells (MEIS1), enteroendocrine cells (CHGA), goblet cells (MUC2), colonocytes (CEACAM7) and infection-specific colonocyte subtype 1 (NDRG1). f) Heatmap showing expression profiles of top differentially expressed genes ranked by change magnitude per cell type per condition. g) Heatmap showing chromatin accessibility patterns of top differentially accessible regions ranked by change magnitude per cell type per condition. h) Bar plot showing number of open chromatin regions with significantly increased (up) or decreased (down) detection rate upon *S*.Tm infection. i) Representative IHC staining of LCN2 in non-infected and *S*.Tm infected colonic epithelium with accumulated mucus (4 days, 5 µl) after 24 h of co-culture. Strong LCN2 accumulation observed at the basal side of goblet cells (magnified insert; black arrows). Scale bar: 100 µm.

**Extended Data Figure 8.**
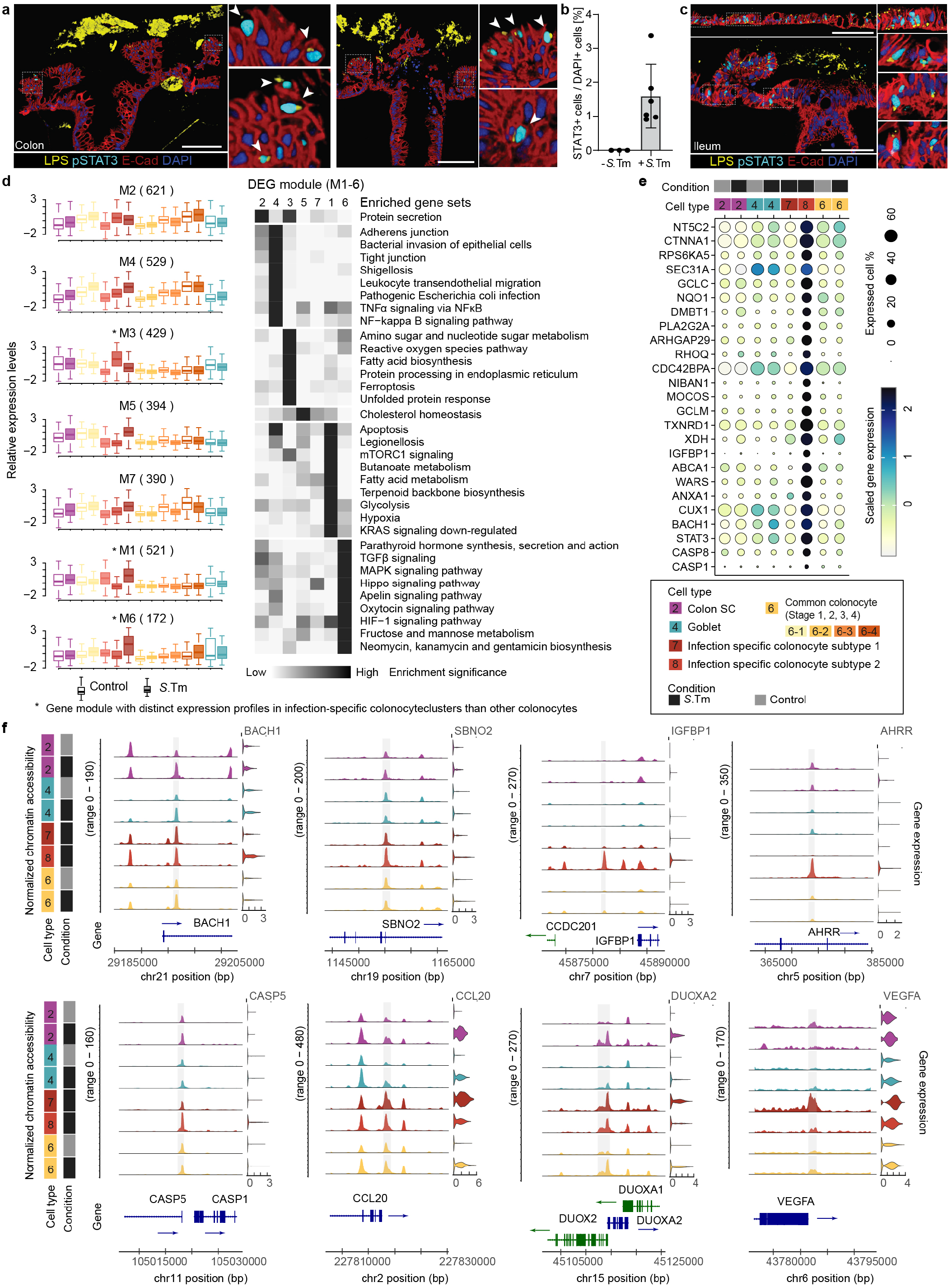
Differential response to *S*.Tm infection across cell types. a) mIF staining of a cross-section of colonic epithelium with accumulated mucus (4 days, 5 µl) and 24 h of *S*.Tm co-culture, showing *S*.Tm (yellow) in the luminal side and areas with epithelium breach and intracellular invasion. STAT3 activation (p-STAT3; cyan) appears in invaded cells, colocalizing with intracellular bacteria (magnified inserts; white arrows). Scale bar: 100 µm. b) Fraction of STAT3+ cells of total DAPI+ cells in cross-sections of non-infected and *S*.Tm infected colonic samples with accumulated mucus (4 days, 5 µl) after 24 h. Bars represent the mean +/-SD. d=1, n=3 (-*S*.Tm), n=6 (+*S*.Tm) (dots). c) mIF staining of a cross-section of ileal epithelium without accumulated mucus infected with *S*.Tm (yellow) for 2 h 30 min before gentamicin wash and 18 h of additional co-culture. Same infection conditions as Extended Data Fig. 6j-k. p-STAT3 (cyan) detected in infected cells. Scale bar: 100 µm. d) Box plots (left) showing distinct expression patterns of infection induced DEGs across cell types between conditions (co-expression gene module index with number of genes in parenthesis) whereby box transparency indicates the condition and box color indicates cell type identity, which follows the color scheme of panel d. Y-axis shows the z-transformed cell type average expression levels of each gene. The five lines of the box represent minimum, 25th percentile, median, 75th percentile, and maximum of each gene module (left). Heatmap showing the scaled KEGG pathway enrichment of each gene module which is defined as the min-max normalized -log10-transformed hypergeometric test nominal P-value (right). e) Dot plot showing distinct transcription program of infection-specific colonocyte subtype 2. f) Normalized expression levels of *S*.Tm infection upregulated genes and the chromatin accessibility levels of their regulatory regions across epithelial cell types in different conditions, highlighting features associated with infection-specific colonocyte subtype 2 (BACH1, SBNO2, IGFBP1, AHRR, CASP5), ubiquitous (CCL20, DUOXA2) and infection-specific colonocyte subtype 1 (VEGFA).

**Extended Data Figure 9.**
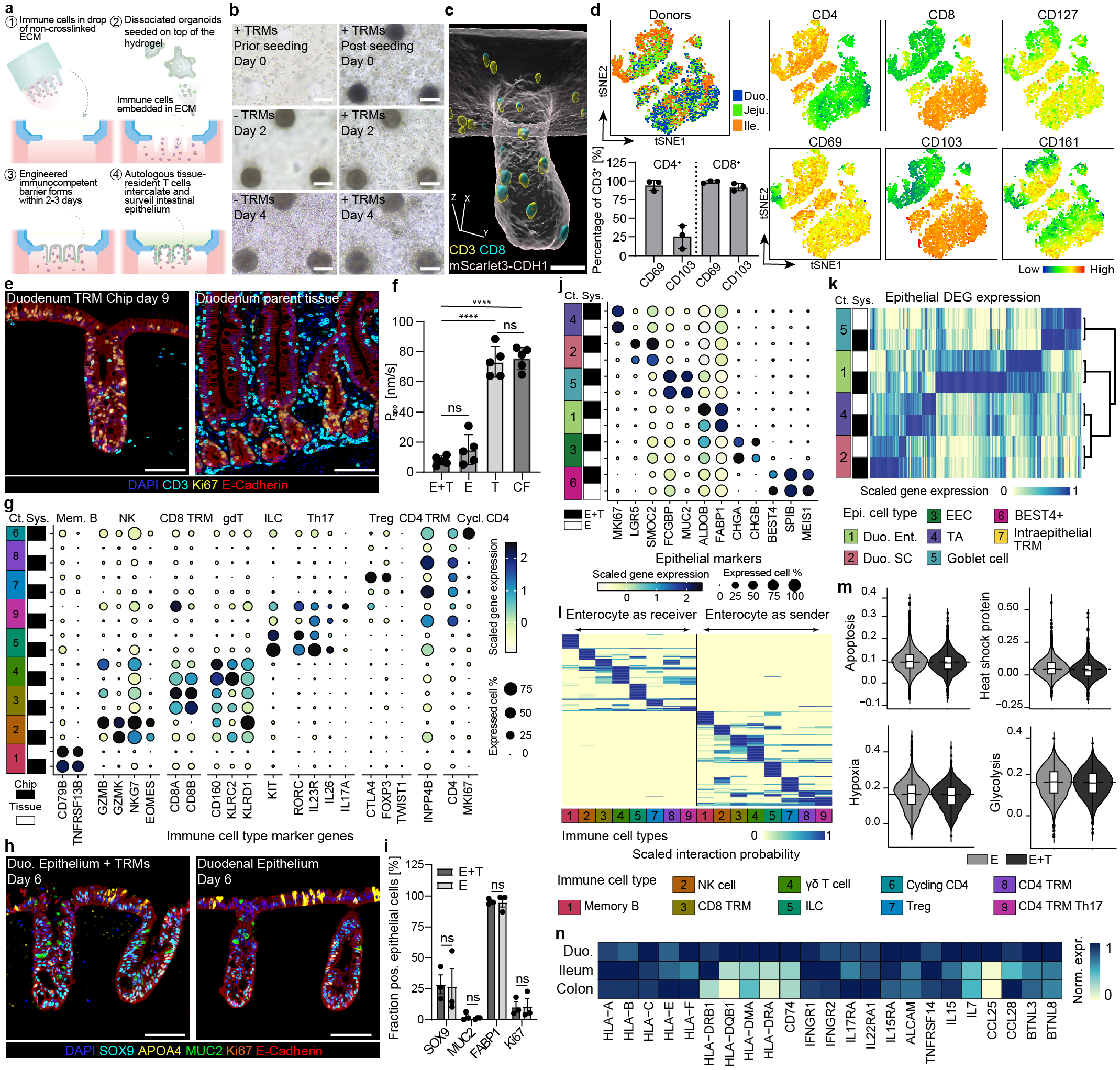
Immune-epithelium co-culture does not differ in canonical marker expression in comparison to epithelium-only engineered intestines. a) Schematic workflow to integrate TRMs into the basal compartment of the engineered intestines. b) Representative regions of engineered duodenal intestines highlighting TRMs co-culture in comparison to epithelium-alone control during early epithelialization. Scale bar: 100 µm. c) Representative 3D-rendered image of a single crypt (view from bottom) with intraepithelial CD3+ (yellow) and CD8+ (cyan) T cells Cas9 CDH1-mScarlet3 reporter-line intestinal epithelium (E-Cadherin, grey) at day 7. Scale bar: 50 µm. d) tSNE embedding on tissue-resident immune cells derived from the small intestine of donors (d=3) at baseline, with cells colored by regions (left) and feature plots showing surface marker expression (right). Barplot highlights the percentage of CD4+ and CD8+ T cells (gated on viable CD45+ CD3+) expressing CD103 and CD69 as mean +/-SD. e) Representative mIF cross-sections of a duodenal crypt at day 9 (Ki67+, yellow) with CD3+ TRMs (cyan) residing in the matrix and epithelial monolayer (E-Cadherin+, red) in comparison to the donor-matched primary tissue. Scale bar: 100 µm. f) Apparent permeability of immunocompetent engineered intestines (E+T) in comparison to epithelium-alone (E), cell-free hydrogel (CF) and TRMs alone (T) at day 3 of culture. Bars reflect the mean of five donors +/-SD (d=5, dots), which were derived from multiple engineered intestines per donor. Unpaired two-tailed t test with Welch’s correction (****P<0.0001, ns = P>0.05). g) Dot plot showing average expression of immune markers of TRMs co-cultured on engineered duodenal intestines for seven days in comparison to reference atlas data. h) Representative mIF image exemplifying patterned canonical marker expression and APOA4+ proximal enterocyte expression between immunocompetent epithelium and epithelium-alone at day 6 of culture. Comparison of canonical epithelial marker expression between the immuno-competent engineered intestines and the intestines cultured in isolation at day 10 of culture. Marker expression normalized to total DAPI+ cells in the epithelium. Bars reflect the mean of three donors (d=3) +/-SD, which were derived from three engineered intestines (n=3) per donor, unpaired two-tailed t test (ns = P>0.05). j) Dot plot showing average expression of canonical intestinal epithelial marker expression of engineered duodenal intestines co-cultured with TRMs and in isolation at day 7 of culture. k) Heatmap showing expression patterns of epithelial DEGs upon TRM coculture across epithelial cell types and conditions at day 7 of culture. l) Heatmap showing scaled CellChat derived interactions probability of directional cell type specific ligand-receptor pairings between enterocytes and TRMs at day 7 of culture. m) Violin plots showcasing Seurat derived module score based on curated lists of genes associated with epithelial cell stress and apoptosis between epithelium co-cultured with TRMs and in isolation at day 7 of culture. n) Comparison of average expression of genes associated to mucosal immune interactions of the intestine across engineered duodenal, ileal, colonic intestines cultured for 10 days. Average expression of each gene was normalized to the maximum expression of each gene per region. Data derived from day 10 scRNAseq engineered tissue atlas (Fig. 1).

**Extended Data Figure 10.**
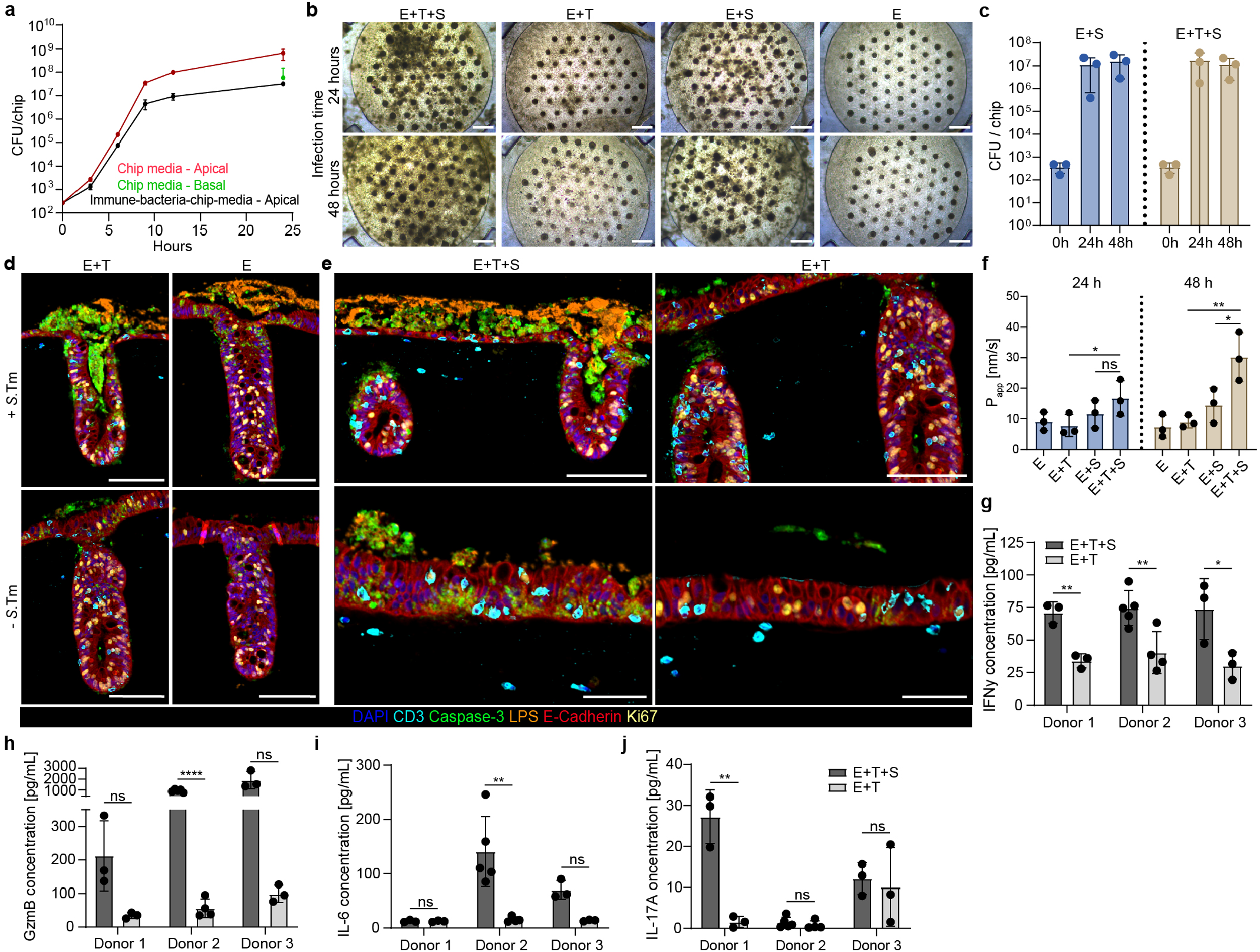
*S*.Tm infection damages epithelium resulting in leaky barrier, epithelial thinning and apoptosis. a) Colony forming units (CFU) of *S*.Tm grown apically in chip-media versus immune-bacteria-chip-media (Suppl. Table 3). The 24 h timepoint includes CFU measurements of the basal compartment at 24 h, where the immune-bacteria-chip-media basal CFU were 0. n=3 averaged +/ SD per medium and timepoint. b) Representative top-view brightfield images of immunocompetent (E+T) and epithelium-only (E) non-infected engineered intestines compared to 24 and 48 hours infected samples (E+T+S, E+S). Scale bar: 500 µm. c) CFU comparison of *S*.Tm in the lumen of the engineered tissue over time between the immunocompetent barrier (E+T+S) and the epithelium co-cultured with *S*.Tm alone (E+S). d-e) Representative mIF images highlighting day-9 duodenal immunocompetent barrier and epithelium-alone infected with *S*.Tm for 48 h (E+T+S, E+S) and the non-infected controls (E, E+T). Showcased images exemplify hallmarks of *S*.Tm infection, namely apoptosis induction (cleaved Caspase-3, green) and barrier thinning (E-Cadherin, red), with CD3+ T cells (cyan) dispersed in the matrix and epithelium. Ki67+ exemplifies patterning of epithelium and few proliferative T lymphocytes (CD3+ Ki67+). Scale bar: 100 µm. Scale bar of magnified images: 50 µm. f) Permeability assessment of engineered immunocompetent and epithelium-only duodenal epithelia at day 8 and day 9, 24 h and 48 h of *S*.Tm post infection, respectively (E+T+S, E+S), compared to non-infected controls (E, E+T). Bars reflect the mean of donors (d=3) +/-SD, which were derived from multiple engineered intestines per donor (d=3). Two-way ANOVA with Tukey’s multiple comparisons test, non-significant (ns = P>0.05) unless otherwise specified (*P_24h_=0.0138, *P_48h_=0.0109, **P_48h_=0.0025). g-j) Cytokines detected in the supernatant of the intestinal barrier after 48 h of infection (nine days of co-culture). Bars reflect engineered intestines (n=3-5; dots) per respective donor (d=3) +/-SD. g) IFNγ secretion, two-tailed unpaired t-test (**P_Don1_=0.0027, **P_Don2_=0.0099, *P_Don3_=0.0420). h) GzmB secretion, two-tailed unpaired t-test with Welch’s correction (ns,P_Don1_=>0.05, ****P_Don2_<0.0001, ns,P_Don3_=>0.05). i) IL-6 secretion, two-tailed unpaired t-test with Welch’s correction (ns,P_Don1_=>0.05, **P_Don2_=0.0067, *P_Don3_=0.0295). j) IL-17A secretion, two-tailed unpaired t-test (**P_Don1_=0.0027, ns,P_Don2_=>0.05, ns,P_Don3_=>0.05).

